# Target enrichment and extensive population sampling help untangle the recent, rapid radiation of *Oenothera* sect. *Calylophus*

**DOI:** 10.1101/2021.02.20.432097

**Authors:** Benjamin J. Cooper, Michael J. Moore, Norman A. Douglas, Warren L. Wagner, Matthew G. Johnson, Rick P. Overson, Angela J. McDonnell, Rachel A. Levin, Robert A. Raguso, Hilda Flores Olvera, Helga Ochoterena, Jeremie B. Fant, Krissa A. Skogen, Norman J. Wickett

## Abstract

*Oenothera* sect. *Calylophus* is a North American group of 13 recognized taxa in the evening primrose family (Onagraceae) with an evolutionary history that may include independent origins of bee pollination, edaphic endemism, and permanent translocation heterozygosity. Like other groups that radiated relatively recently and rapidly, taxon boundaries within *Oenothera* sect. *Calylophus* have remained challenging to circumscribe. In this study, we used target enrichment, flanking non-coding regions, summary coalescent methods, tests for gene flow modified for target-enrichment data, and morphometric analysis to reconstruct phylogenetic hypotheses, evaluate current taxon circumscriptions, and examine character evolution in *Oenothera* sect. *Calylophus*. Because sect. *Calylophus* comprises a clade with a relatively restricted geographic range, we were able to extensively sample across the range of geographic and morphological diversity in the group. We found that the combination of exons and flanking non-coding regions led to improved support for species relationships. We reconstructed potential hybrid origins of some accessions and note that if processes such as hybridization are not taken into account, the number of inferred evolutionary transitions may be artificially inflated. We recovered strong evidence for multiple origins of the evolution of bee pollination from ancestral hawkmoth pollination, the evolution of edaphic specialization on gypsum, and permanent translocation heterozygosity. This study applies newly emerging techniques alongside dense infraspecific sampling and morphological analyses to effectively address a relatively common but recalcitrant problem in evolutionary biology.

## INTRODUCTION

The challenges of reconstructing species histories for groups that arose through recent, rapid radiations are well established. Phylogenetic signal can be obscured by processes such as incomplete lineage sorting (ILS) and gene flow (Maddison and Knowles 2006; Knowles and Chan 2008; Christie and Knowles 2015), resulting in short branch lengths and conflicting gene tree topologies. Consequently, approaches that use few loci or concatenation may fail to resolve the most accurate species tree (Eckert and Carstens 2008; Leaché et al. 2014; Xi et al. 2014; Giarla and Esselstyn 2015). This may be particularly common in plants that are thought to have experienced rapid or recent radiation with ongoing hybridization and high levels of ILS. The application of target enrichment methods that efficiently sequence hundreds of nuclear loci, coalescent-based phylogenetic methods that account for ILS and gene flow, and extensive sampling of morphologically diverse populations across the geographic range should allow for more accurate representations of phylogenetic relationships (Maddison and Knowles 2006; Knowles and Chan 2008; Knowles 2009; Mamanova et al. 2010; Lemmon et al. 2012; Straub et al. 2012; Bryson et al. 2014; Weitemier et al. 2014; Mandel et al. 2014; Stephens et al. 2015; Johnson et al. 2016).

*Oenothera* sect. *Calylophus* currently comprises seven species (thirteen taxa) with a center of diversity in western Texas, southern New Mexico, and north-central Mexico (Fig. 1; Towner 1977; Turner and Moore 2014; Wagner 2021). Previous analyses suggest that *Oenothera* sect. *Calylophus* forms a well-supported, morphologically coherent clade with a relatively restricted geographic range (Towner 1977; Levin et al. 2004; Wagner et al. 2007; Turner and Moore 2014; Wagner 2021). However, as with other groups that have experienced rapid radiations, taxon boundaries within *Oenothera* sect. *Calylophus* have been challenging to define, likely due to phenomena such as overlapping morphological boundaries, ongoing introgression, and incomplete lineage sorting.

**Figure 1.**
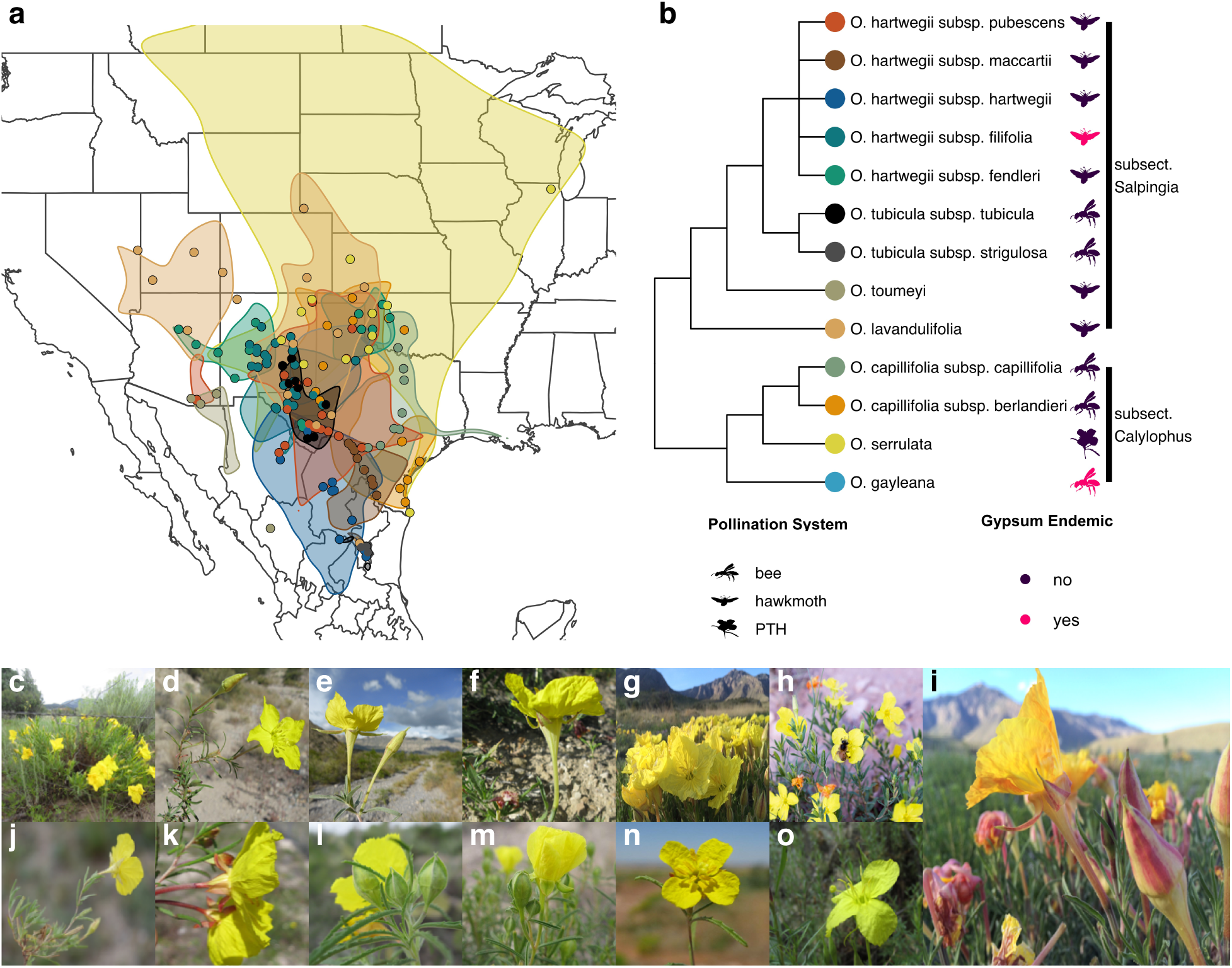
(a) Range map of all taxa in *Oenothera* sect. *Calylophus* (based on Towner 1977). Sampling locations of leaf tissue samples (points; color corresponds to taxa in cladogram to the right [Figure 1b]) and estimated taxon ranges (polygons; colors correspond to Figure 1b) proposed by Towner (1977) and Turner and Moore (2014). (b) Estimated cladogram of *Oenothera* sect. *Calylophus* sensu Towner (1977) and Turner and Moore (2014). Symbols to the right of tip labels signify pollination system (bee, hawkmoth, or Permanent Translocation Heterozygosity [PTH]) and the symbol color specifies whether a given taxon is a gypsum endemic (purple = no, pink = yes). Photo panels: (c) *Oenothera hartwegii* subsp. *fendleri* (d) *Oenothera hartwegii* subsp. *filifolia* (e) *Oenothera hartwegii* subsp. *hartwegii* (f) *Oenothera hartwegii* subsp. *maccartii* (g) *Oenothera hartwegii* subsp. *pubescens* (h) *Oenothera tubicula* subsp. *tubicula* (i) *Oenothera lavandulifolia* (j) *Oenothera tubicula* subsp. *strigulosa* (k) *Oenothera capillifolia* subsp. *berlandieri* (l) *Oenothera capillifolia* subsp. *capillifolia* (m) *Oenothera gayleana* (n) *Oenothera serrulata* (o) *Oenothera toumeyi*

In the most comprehensive study of the group to date, Towner (1977) circumscribed taxa using morphology, breeding system, geography, and ecology, but it was noted (and our field observations confirm) that characters often overlap among taxa (Towner 1977). Taxa within *Oenothera* sect. *Calylophus* are divided into two easily recognizable subsections: subsect. *Salpingia* and subsect. *Calylophus* (Towner 1977; Wagner et al. 2007). Pollination varies between the two subsections; flowers of subsect. *Salpingia* are adapted to vespertine pollination by hawkmoths, except for *O. tubicula*, which opens in the morning and is primarily pollinated by bees (Towner 1977), while taxa in subsect. *Calylophus* are predominantly bee-pollinated (Towner 1977). Taxa in both subsections have geographic ranges that partially (or even largely) overlap, resulting in occasional morphologically intermediate populations (Towner 1977). Although confounding for morphological-based analyses, this observed pattern of reticulation is consistent with a recent, rapid radiation occurring in parallel with climatic fluctuations and increasing aridity in the region since the Pleistocene (Raven 1964; Towner 1977; Nason et al. 2002; Katinas et al. 2004). Hawkmoth pollination, which is ancestral in Onagraceae and common in *Oenothera* sect. *Calylophus*, is known to result in long-distance pollen movement (Stockhouse 1973; Skogen et al. 2019). Therefore, gene flow may have been extensive over the evolutionary history of hawkmoth-pollinated taxa, increasing the chances that processes such as historical introgression may obscure phylogenetic signal in extant plants (Elrich and Raven 1969). With a phylogenomic approach that samples hundreds of nuclear loci, we may better illuminate both the history of these species and the key evolutionary processes related to speciation in this group.

Understanding speciation in angiosperms remains a fundamental question in evolutionary biology (Barrett et al. 1996; Rajakaruna 2004; van der Niet et al. 2006; Wilson et al. 2007; Crepet and Niklas 2009; Peakall et al. 2010; Xu et al. 2011; Van der Niet and Johnson 2012; Boberg et al. 2014). *Oenothera* section *Calylophus* has an evolutionary history that likely involves changes in reproductive system (pollinator functional group, breeding system) and edaphic endemism. For example, there are thought to be two independent shifts between pollinators from hawkmoth to bee pollination (Towner 1977; Fig. 1b), despite many studies in other plant groups showing a directional bias in shifts from bee to hummingbird or hawkmoth pollination (Barrett et al. 1996; Wilson et al. 2007; Thomson and Wilson 2008; Tripp and Manos 2008; Barrett 2013). However, pollinator shifts that do not follow this sequence may be more likely when the extent of trait divergence and specialization does not completely inhibit secondary pollinators such as bees, as has been suggested in *Oenothera* sect. *Calylophus* (Stebbins 1970; Tripp and Manos 2008; Van Der Niet et al. 2014). Shifts to autogamy are also frequent across angiosperms and in Onagraceae alone there are an estimated 353 shifts to modal autogamy (i.e. mostly autogamous; Raven 1979). *Oenothera* sect. *Calylophus* also includes at least one autogamous species, *O. serrulata*, which exhibits permanent translocation heterozygosity, a phenomenon in which all chromosomes are translocationally heterozygous (PTH; Towner 1977) (Fig. 1b). While the evolution of PTH has been assessed in molecular phylogenetic analyses across Onagraceae (Johnson et al. 2009; Hollister et al. 2019), no study to date has examined this transition in a well-sampled clade with extensive population sampling. Lastly, abiotic ecological factors such as edaphic specialization are also known to drive speciation in some groups (Rajakaruna 2004; van der Niet et al. 2006), including in *Oenothera* sect. *Pachylophus* (Patsis et al. in press). For example, serpentine endemics represent ∼10% of the endemic flora in California even though serpentine soils account for about 1% of terrestrial habitat in the state (Brady et al. 2005). Similarly, the Chihuahuan Desert is comprised of numerous isolated islands of gypsum outcrops and current estimates suggest that at least 235 taxa from 36 different plant families are gypsum endemics (Moore and Jansen 2007; Moore et al. 2014). It is suspected that gypsum endemism has also evolved independently in *Oenothera* sect. *Calylophus* at least twice (Towner 1977; Turner and Moore 2014; Fig. 1b). Ultimately, to understand the role that these transitions have played in shaping the diversity of *Oenothera* sect. *Calylophus*, a robust phylogeny is required.

Here, we use target enrichment, summary coalescent methods, and morphometric analyses to reconstruct a phylogenetic relationships, leveraging this information to re-examine previous taxonomic concepts, test for instances of hybridization, and resolve the history of pollinator shifts, PTH, and gypsum endemism in *Oenothera* sect. *Calylophus*. Target enrichment is a cost-effective method for sequencing hundreds of loci across a high volume of samples, producing highly informative datasets for phylogenetics (Lemmon et al. 2012; Straub et al. 2012; Mandel et al. 2014; Weitemier et al. 2014; Heyduk et al. 2015; Stephens et al. 2015; Johnson et al. 2016). While target enrichment is generally designed to capture coding regions, a significant proportion of flanking non-coding regions can be recovered (the “splash-zone”; Weitemier et al. 2014). The inclusion of non-coding regions may be particularly informative for recent radiations, since these regions are less constrained by selective pressures and may contain on average more informative sites at shallower time scales (Folk et al. 2015). We included these flanking non-coding regions in our sequence alignments to evaluate their impact on reconstructing lower-order relationships. Importantly, we sampled extensively, including individuals from numerous populations across the geographic and morphological ranges of all thirteen taxa in the section (Fig. 1a). This study presents an example of how combining these molecular techniques with dense sampling and morphological analysis can be used to effectively address a common problem in evolutionary biology.

## MATERIALS AND METHODS

A total of 194 individuals spanning the geographic, morphological, and ecological ranges of all 13 recognized taxa in *Oenothera* sect. *Calylophus* [following Towner (1977) and Turner and Moore (2014)] were included in this study (Fig. 1a, S1, S2) along with 8 outgroups representing other major sections of *Oenothera* (*Eremia, Gaura, Kneiffia, Lavauxia, Oenothera, Pachylophus*, and *Ravenia*) and other genera (*Chylismia* and *Eulobus*) in Onagraceae (S1, S2). DNA was extracted from fresh, silica-dried leaf tissue (S3). PTH status was confirmed or reassessed for individuals in subsect. *Calylophus* by assessing pollen fertility, when flowers were present, using a modified Alexander stain (Alexander 1969, 1980; S3).

Target nuclear loci for enrichment were determined by clustering transcriptome assemblies of *Oenothera serrulata* (1KP accession *SJAN*) and *Oenothera capillifolia* subsp. *berlandieri* (1KP accession *EQYT*). Starting with the 956 phylogenetically informative *Arabidopsis* loci identified by Duarte et al. (2010; S3), we identified 322 homologous, single-copy loci in our clusters and used these in the probe design process. Libraries were enriched for these loci using the MyBaits protocol (Arbor Biosciences, Ann Arbor, MI, USA) and sequenced on an Illumina MiSeq (2 x 300 cycles, v3 chemistry; Illumina, Inc., San Diego, California, USA). Raw reads have been deposited at the NCBI Sequence Read Archive (BioProject PRJNA544074; See S3 for details). Reads were trimmed using Trimmomatic (Bolger et al. 2014; S3) and trimmed, quality-filtered reads were assembled using HybPiper (Johnson et al. 2016). From the assembled loci, we produced two datasets: **“exons”,** constiting of exon-only alignments, and **“supercontig”,** consisting of alignments containing both the exon alignment and flanking non-coding regions (the “splash-zone” per Weitemier 2014 and reconstructed using supercontigs produced by HybPiper). We used these two datasets to test the most recent taxonomic circumscription of the group with several methods: (1) phylogenetic inference of concatenated alignments (two analyses: exons and supercontigs) using RAxML (Stamatakis 2014), (2) ASTRAL-II (Mirarab and Warnow 2015; Sayyari and Mirarab 2016) species tree inference (two analyses: exons and supercontigs), (3) SVD Quartets (Chifman and Kubatko 2014, 2015) species tree inference (one analysis: supercontigs), (4) Phyparts (Smith et al. 2015; one analysis: supercontigs), (5) IQtree (Minh et al. 2018) with both gene and site concordance factors (one analysis: supercontigs).

We used HyDe (Blischak et al. 2018) to test for putative hybrid origins of selected taxa and accessions by calculating D-Statistics (Green et al. 2010) for a set of hypotheses (S3). To further characterize population-level processes or genetic structure within sect. *Calylophus*, we extracted and filtered SNPs by mapping individual reads against reference supercontigs (see https://github.com/lindsawi/HybSeq-SNP-Extraction) and used Discriminant Analysis of Principal Components (Jombart et al. 2010) as implemented in the R package *adegenet* (Jombart 2008) and the snmf function in the LEA package (Frichot and François 2015) in R (R Core Team, 2020; S3).

We evaluated current taxonomic concepts and patterns of morphological variation by measuring character states for vegetative and floral structures that have been used historically to discriminate taxa in *Oenothera* sect. *Calylophus* (Towner 1977): plant height, leaf length (distal), leaf width (distal), leaf length/width ratio (distal), leaf length (basal), leaf width (basal), leaf length/width (basal), sepal length, and sepal tip length (S3). Measurements were made for 125 of the sequenced samples (S11); we were unable to measure all traits for 73 samples because we did not have access to the herbarium vouchers, or the trait of interest was not captured on the voucher, therefore some samples were dropped from the analysis due to missing values. Finally, the number of transitions and inferred ancestral conditions of reproductive system were mapped onto an ASTRAL species tree, with individuals grouped into species, using the stochastic mapping function in the R package *phangorn* version 2.5.5 (Schliep 2011; S3).

## RESULTS AND DISCUSSION

### Sequencing and Phylogenetic Results

Sequencing resulted in a total of 80,273,296 pairs of 300-bp reads with an average of 625,323 reads per sample. Following quality filtering, assembly and alignment, we recovered 204 loci that were present in at least 70% of the samples. Across all datasets and analyses, *Oenothera* sect. *Calylophus* was monophyletic. At the subsection level there was strong agreement in topology between concatenation and coalescent-based trees. For example, subsect. *Calylophus* was recovered as sister to *Oenothera* subsect. *Salpingia* (minus *O. toumeyi*) with strong support across all analyses and *O. toumeyi* [considered by Towner (1977) to be in subsect. *Salpingia*; Fig. 2, S4-7] was recovered as sister to subsect. *Calylophus* across all trees, with strong bootstrap support. Within subsect. *Calylophus* there was poor resolution for currently recognized taxa in all analyses, whereas taxon relationships were better resolved in subsect. *Salpingia* (Fig. 2, S4-7). With coalescent-based tree reconstruction, most taxa *sensu* Towner (1977) were recovered as monophyletic with moderate to strong support (Fig. 2, S6-8). In contrast, both the exon and supercontig concatenation trees recovered most currently recognized taxa as non-monophyletic (S4, S5). Given that concatenation has been shown to produce incorrect topologies in the presence of high ILS (Roch and Steel 2015) and that *Oenothera* sect. *Calylophus* underwent recent radiation, we believe the paraphyly of taxa in both concatenation trees might be artifactual. We therefore interpret relationships based on our coalescent-based trees, which comprise the focus for the remainder of the paper (Fig. 2, S6-8).

**Figure 2.**
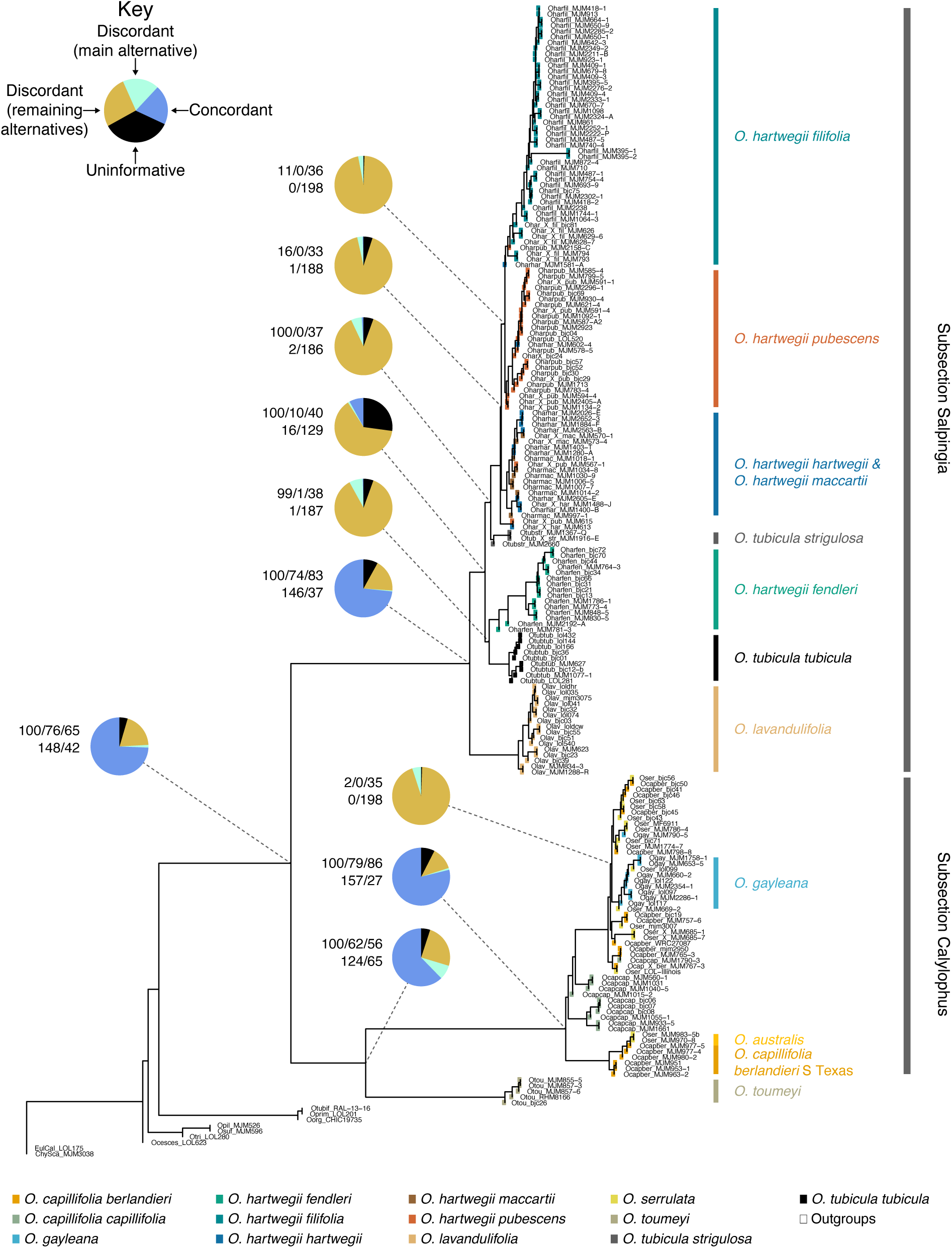
ASTRAL-II summary coalescent tree constructed using the “Supercontig” dataset with 100 bootstraps. At relavent nodes, piecharts represent Phyparts analysis (blue = concordant, green = most frequent conflict, yellow = all other conflict, black = uninformative gene trees), top row of support values are bootstrap values from ASTRAL-II, and gCF and sCF from IQTree (BS/gCF/sCF), bottom two support values are number of concordant gene trees for the node and total number of gene trees minus the number of concordant gene trees at that node (concord/discord). Colored tip points correspond to taxon designation.

To understand whether summary coalescent relationships display a consistent signal across the genome, we quantified gene tree and site concordance using Phyparts and IQtree. We found that gene tree concordance was highest at the deepest nodes at the species level where we expected less ILS and more time between speciation events (Fig. 2, S8, S9). Correspondingly, gene tree concordance was lowest at the subspecies level where increased sharing of ancestral alleles and ongoing gene flow are more likely (Fig. 2, S8, S9). For example, within *O. hartwegii* less than 1% of genes were concordant for bifurcations representing all currently recognized taxa at the subspecies level (Fig. 2, S8, S9). For species-level nodes with high support and high gene tree concordance, site concordance was also high; for example, *O. lavandulifolia* had high bootstrap support in summary coalescent trees (BS = 100), high gene tree concordance (Phyparts = 94% concordance, gCF = 74), and high site concordance (sCF = 83; Fig. 2, S9h). For subspecies with high support, but low gene tree concordance, site concordance was moderate. For example *O. hartwegii* subsp. *fendleri* had high bootstrap support (BS = 99), low gene tree concordance (Phyparts = <1% concordance, gCF = 1), and moderate site concordance (sCF = 38; Fig. 2, S9b). For subspecies that were monophyletic in our coalescent-based trees, but that had low support and low gene tree concordance, site concordance was moderate with an average of 35% of sites in agreement for taxa at these nodes (S9[c,f,g,v]). For example *O. hartwegii* subsp. *filifolia* had low bootstrap support, low gene tree concordance (Phyparts = <1% concordance, gCF = 0), and moderate site concordance (sCF = 36; Fig. 2, S9c). This is an important finding because while species tree relationships can be obscured by ILS, site concordance factors, which may be less constrained and less subject to ILS at shallower evolutionary timescales, provide a key alternative method of support (Minh et al. 2018).

In general, topologies of exon and supercontig datasets were similar, with no major differences in clade membership, but the inclusion of the flanking non-coding regions increased support at shallow nodes in our trees. However, this trend was not universal. For example, in subsect. *Salpingia*, using the supercontig dataset decreased support slightly for one taxon (*O. hartwegii* subsp. *filifolia*), and in subsect. *Calylophus* it led to paraphyly of another (*O. capillifolia* subsp. *capillifolia*). For six other taxa, our results showed that using supercontigs increased bootstrap support. Therefore, these results demonstrated a net benefit of including flanking non-coding regions for resolving relationships among closely related taxa.

### Hybridization and Geneflow

Using concatenated loci from the supercontig dataset, we used HyDe (Blischak et al. 2018) to test for signals of hybridization. We used 552,521 sites and tested 22 hypotheses for either individuals or groups suspected to be of hybrid origin based on field observations of morphological intermediacy, geographic location, and topological position in our coalescent-based trees and found evidence of hybridization in three individuals representing two taxa, both in subsect. *Salpingia*. The highest signal of hybridization, with a gamma value (γ^ ) of 0.947 suggesting more historic gene-flow, was observed in one of three sampled individuals of *O. tubicula* subsp. *strigulosa* (*MJM1916.E*). This involved admixture between *O. tubicula* subsp. *tubicula* and the clade consisting of *O. hartwegii* subsp*. hartwegii* and *O. hartwegii* subsp. *maccartii* (Z-score = 5.585, p-value = 0.000, γ^ = 0.947; Fig. 3a, S10). We also detected significant levels of hybridization, with γ^ ranging from 0.332 to 0.338 suggesting more contemporary gene-flow, in two individuals in *O. hartwegii* subsp. *pubescens*, *BJC29* (Z-score = 2.378, p-value = 0.009, γ^ = 0.338) and *MJM594* (Z-score = 2.094, p-value = 0.018, γ^ = 0.332). This more recent gene flow involved admixture between *O. hartwegii* subsp. *pubescens* and the clade consisting of *O. hartwegii* subsp. *hartwegii* and *O. hartwegii* subsp. *maccartii* (Fig. 3a, S10).

**Figure 3.**
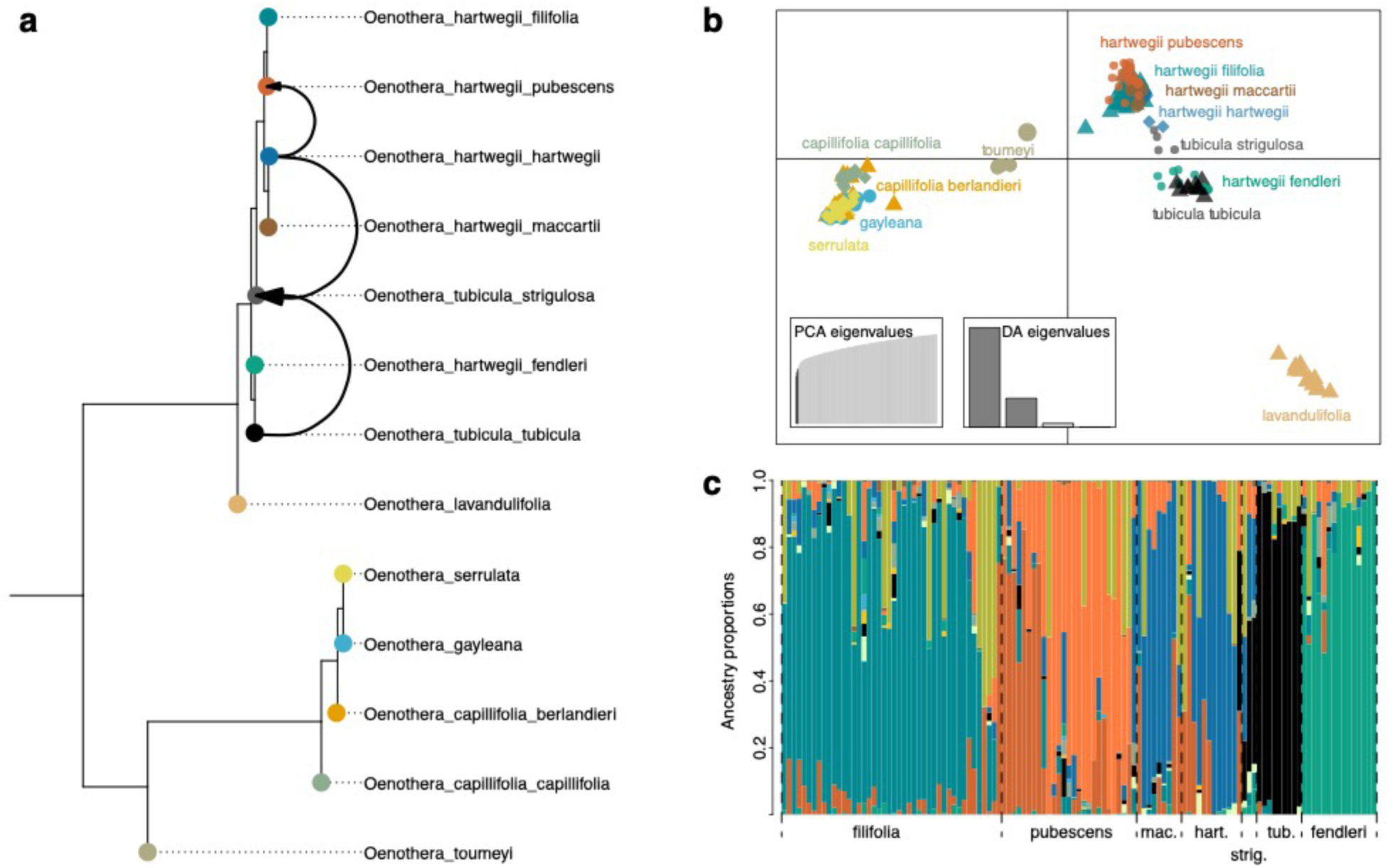
(a) Summary of HyDe Analysis annotated on ASTRAL-III species tree constructed using the “Supercontig” dataset; black arrows represent direction of admixture detected by HyDe anaylsis. (b) Discriminant Analysis of Principal Components based on a filtered set of SNPs extracted from the entire supercontig dataset, and (c) sNMF plot of inferred ancestry coefficients using the same set of filtered SNPs but limited to subsection *Salpingia* (minus *O. lavandulifolia*)

The finding that one individual of *O. tubicula* subsp. *strigulosa* may be of hybrid origin is consistent with gene flow between *O. tubicula* subsp. *strigulosa* and its sister taxon *O. tubicula* subsp. *tubicula* (sensu Towner 1977). In our coalescent-based analyses, the two subspecies of *O. tubicula* were not recovered as sister taxa, and this relationship was strongly supported (S9[j-k]). If the two *O. tubicula* taxa arose independently, this would support the hypothesis that bee pollination arose in *Oenothera* sect. *Calylophus* independently three times. However, while the summary coalescent analyses we utilized to estimate phylogenies accounted for ILS in tree estimation, they did not account for gene flow (Meng and Kubatko 2009; Gerard et al. 2011; Kubatko and Chifman 2019). Our HyDe results may support the hypothesis that *O. tubicula* subsp. *strigulosa* has experienced gene flow from two closely related taxa, and may have hybrid origins resulting from crossing between *O. tubicula* subsp. *tubicula* and *O. hartwegii* subsp. *hartwegii* (Fig. 3a, S10). This is consistent with Towner’s interpretation of this taxon, which he hypothesized may represent a stabilized derivative of introgression between *O. tubicula* subsp. *tubicula* and *O. hartwegii* subsp. *hartwegii* (Towner 1977). Therefore, the placement of *O. tubicula* subsp. *strigulosa* as sister to the rest of the *O. hartwegii* species complex in our trees may result from past gene flow and hence may not represent independent origins of bee pollination in subsect. *Salpingia*. These results underscore the importance of explicitly including tests for hybridization in phylogenetic studies. In the case of these data, estimating a species tree given a set of gene trees within a coalescent framework without considering other non-ILS sources of signal conflict could artificially inflate the number of inferred evolutionary transitions.

Our HyDe results also suggest that at least some of the morphological intermediacy and overlap among taxa in the group is due to continued, or at least recent, gene flow. For example, both *O. hartwegii* subsp. *pubescens* individuals that are inferred to have significant levels of admixture were collected from morphologically intermediate populations of *O. hartwegii* subsp. *pubescens* and *O. hartwegii* subsp. *hartwegii*. In addition, *O. hartwegii* subsp. *hartwegii* was a parent in all three hybridization events (S10). Thus, gene flow may explain this taxon’s non-monophyly in our summary coalescent results. However, despite the often confounding patterns of overlapping morphological variation among closely related taxa in subsect. *Salpingia*, this pattern does not necessarily appear to be the result of admixture, as many of the tests for hybridization based on field observations were not significant (Fig 3a, S10). What is also clear from these results is that much like collecting hundreds of nuclear genes provides a more nuanced picture of phylogenetic signal and taxon relationships, our results show that collecting multiple individuals from across the geographic and morphological ranges is necessary for a more complete picture of relationships among closely related taxa.

After filtering, we extracted a set of 9,728 single nucleotide polymorphisms (SNPs) from both coding and non-coding regions. A Discriminant Analysis of Principal Components (DAPC; Fig. 3b) using these data clearly distinguishes *Oenothera* subsect. *Salpingia* from *O.* subsect. *Calylophus*, with *O. toumeyi* intermediate between the two, which is consistent with the phylogenetic results presented here. Additionally, the DAPC identifies *O. lavandulifolia* as a distinct genetic cluster from the remaining taxa in subsection *Salpingia*. The overlap between taxa, for example between the remaining taxa in subsection *Salpingia*, is consistent with the high levels of gene tree discordance identified by PhyParts (Fig. 3a). For this latter group of taxa, we computed estimates of ancestry coefficients using snmf, which suggests a substantial amount of shared ancestral polymorphisms while also showing some evidence of clear genetic structure among taxa (Fig. 3c). Consistent with the phylogenetic analyses, there does not appear to be any clear genetic distinction between *O. hartwegii* subsp. *hartwegii* and *O. hartwegii* subsp. *maccartii*, whereas *O. hartwegii* subsp. *fendleri*, *O. tubicula* subsp. *tubicula,* and *O. hartwegii* subsp. *filifolia* appear to be largely distinct.

### Morphological Analysis

We conducted morphometric Principal Components Analysis (PCA) to determine if morphological patterns were consistent with phylogentic results and to examine if specific characters could be used to diagnose taxa as circumscribed by our phylogenetic analysis. Towner (1977) observed overlapping and confounding patterns of morphological variation among taxa within subsections, particularly within the *O. hartwegii* species complex. Despite this, because some taxa (e.g., *O. hartwegii* subsp. *fendleri*) were strongly supported by our summary coalescent trees we expected that they would be well distinguished in morphometric analysis.

The main traits that separated taxa in subsect. *Salpingia* were leaf traits and plant size, while in subsect. *Calylophus* the main traits that separated taxa were sepal traits. In subsect. *Salpingia*, PC1 accounted for 43.1% of variance in PCA, while PC2 accounted for 28.5% (Fig. 4b). Morphological characters most associated with PC1 were leaf width (distal and basal), plant height, and leaf length/width ratio (distal and basal). Those associated with PC2 were leaf length (distal and basal), sepal length, and sepal tip length (Fig. 4b). In subsect. *Calylophus*, PC1 accounted for 44.4% of explained variance and PC2 accounted for 26.2% (Fig. 4c). The characters most associated with PC1 in subsect. *Calylophus* include leaf length (distal and basal), sepal length, and sepal tip length. Those most associated with PC2 were leaf length/width ratio (distal and basal) and distal leaf width (Fig. 4c).

**Figure 4.**
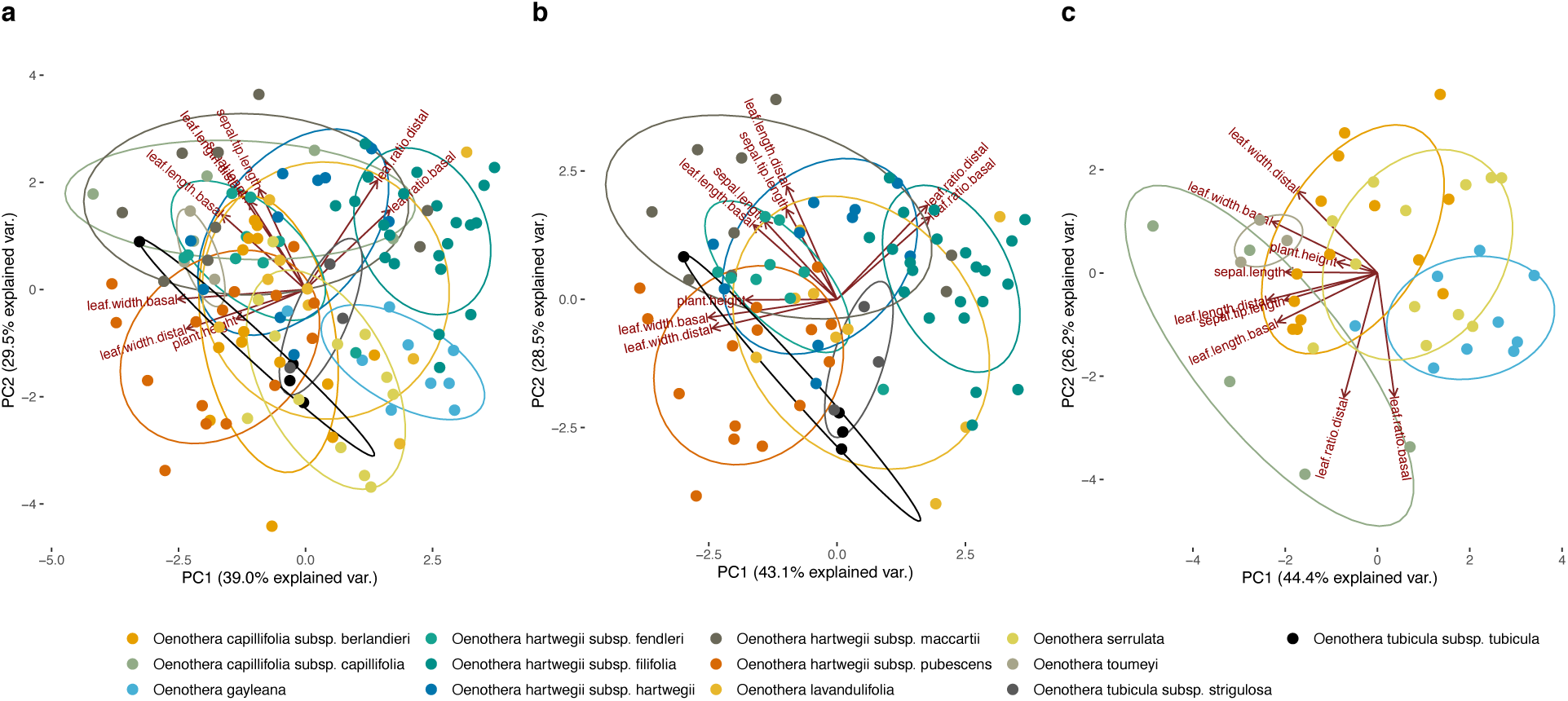
Morphometric Principal Components Analysis (PCA) using 9 morphological characters (plant height, leaf length [basal and distal], leaf width [basal and distal], leaf length/width ratio (basal and distal), and sepal tip length) for (a) Section *Calylophus* (b) Subsection *Salpingia* without *O. toumeyi*, and (c) Subsection *Calylohus* with *O. toumeyi* included.

Our results support Towner’s understanding of taxon boundaries by underscoring previous difficulties in identifying individuals in this group based on morphology (Towner 1977). We found substantial overlapping morphological variation among currently recognized taxa in both subsections, though some taxa exhibited better grouping than others. The amount of overlap between taxa was not a function of the strength of tree support for a given taxon in our summary coalescent results. For example, *O. hartwegii* subsp. *fendleri*, a taxon that was well supported in our summary coalescent trees, exhibited some of the highest degree of overlap with other taxa in PCA space. Conversely, both *O. hartwegii* subsp. *filifolia* and *O. hartwegii* subsp. *pubescens*, two taxa that formed poorly supported clades in our trees, formed clusters on the outer edges of PCA space and had less overlap than other taxa (Fig. 4b). Interestingly, *O. hartwegii* subsp. *hartwegii*, the taxon that was identified as a parent in all three instances of admixture in our HyDe analysis, also overpas morphologically with most other taxa in subsect. *Salpingia* (Fig. 4b). This is not surprising given that it is widely distributed in northern Mexico and western Texas and frequently comes into contact with related taxa resulting in sympatric populations and occasional morphologically intermediate populations.

### Implications for Reproductive Systems and Edaphic Endemism

Our results show that shifts from hawkmoth to bee pollination likely occurred twice in sect. *Calylophus* (S3) and thus may be more common in *Oenothera* than previously thought. The strongly supported sister relationship of *O. toumeyi* to remaining subsect. *Calylophus* in our summary coalescent results is consistent with two independent shifts to bee pollination, once in the ancestor of subsect. *Calylophus*, and another in subsect. *Salpingia* on the branch leading to *O. tubicula* (Fig. 2). Independent shifts to bee pollination from hawkmoth pollination are perhaps not surprising considering that within sect. *Calyophus*, hawkmoth-pollinated floral forms exhibit variation between populations in hypanthium length and diameter and do not prevent occasional pollination by bees (Lewis 2015; Towner 1977). Hawkmoth-pollinated taxa in sect *Calylophus* exhibit vespertine anthesis, which separates them temporally from diurnal bees, but variation in the timing of anthesis is also common between populations (Towner 1977), and hawkmoths are documented to vary greatly in abundance spatiotemporally (Miller 1981; Campbell et al. 1997; Artz et al. 2010). In cases of pollen limitation, night-blooming plants benefit from bimodal pollination between moths and bees by acquiring pollinator assurance against yearly variation or local extinction of specific pollinators, as shown for other *Oenothera* species (Barthell and Knops 1997; Artz et al. 2010), *Lonicera japonica* (Miyake and Yahara 1998) and for night-blooming *Ancistrophora* cacti (Schlumpberger et al. 2009).

Additionally, it has been shown that florivore-mediated selection drives floral trait shifts in sect. *Calylophus* towards bee-pollinated floral forms (Jogesh et al. 2017; Bruzzese et al. 2019). Variation in reproductive traits that allows some continued pollination by bees provides an alternative mode of pollen transfer and may represent a mechanism for ensuring pollination. While studies have shown that premating barriers contribute greatly to reproductive isolation (Stanton et al. 2016), our results show that multiple, indepdendent shifts from hawkmoth to bee pollination and associated morphological changes, such as the shorter floral tube length of bee pollinated flowers, may occur in sect. *Calylophus*, and hence may not be a particularly reliable character for diagnosing taxa in this group.

Stochastic mapping (supplemental) suggests that there are multiple origins of permanent translocation heterozygosity (PTH) in sect. *Calylophus* While ring chromosomes are common and found in all taxa in sect. *Calylophus*, PTH is currently known from only one taxon, *O. serrulata*. Because neighboring populations of *O. serrulata* and its putative progenitor O*. capillifolia* subsp. *berlandieri* often resemble each other phenetically, Towner (1977) hypothesized that *O. serrulata* may have originated multiple times through independent origins of translocation heterozygosity in different geographic regions, and may be best recognized as “a complex assemblage of populations having a common breeding system.” However, this has never before been explored in a phylogenetic context, nor has it been clearly demonstrated with phylogenetic studies in Onagraceae. In our summary coalescent trees, all currently recognized taxa in subsect. *Calylophus* were paraphyletic and *O. serrulata* was scattered throughout the subsection (Fig. 2, S6, S7). Although support values are not always high for the positions of various individuals of *O. serrulata*, there is at least one well defined, well supported split among populations of *O. serrulata*. In our summary coalescent trees, the two *O. serrulata* accessions from south Texas (*MJM970* & *MJM983*) grouped with other south Texas populations of *O. capillifolia* subsp. *berlandieri* with generally strong support (Fig. 2, S6, S7). This relationship was supported in PCA space as well, where *MJM983* was morphologically more similar to the south Texas *O. capillifolia* subsp. *berlandieri* accessions than to other *O. serrulata* (Fig 4C). Our results are therefore consistent with an independent origin of PTH in coastal Texas populations of *O. serrulata*, demonstrating at minimum two origins of PTH (see *Taxonomic Implications* below). However we cannot rule out other independent origins of PTH. Given the prevalence of translocations among partial sets (i.e. not all seven) of homologous chromosomes in *O.* section *Calylophus* (Towner 1977), and if this could be considered an intermediate step towards “complete” PTH, perhaps it is not surprising to reconstruct multiple origins. Our results suggest that a more complete assessment of the extent and distribution of this phenomenon in section *Calylophus* is needed.

Independent origins of gypsum endemism in sect. *Calylophus* are also supported by our analyses (supplemental). Edaphic specialization is a fundamental driver of speciation in plants and contributes greatly to endemism and species diversity in areas with geologically distinct substrates such as gypsum and serpentine outcrops (Kruckeberg 1984; Anacker et al. 2011; Cacho and Strauss 2014; Moore et al. 2014). To date, two gypsum endemic taxa have been described in sect. *Calylophus*, one in each subsection: *O. hartwegii* subsp. *filifolia*, which is relatively widespread on gypsum in New Mexico and trans-Pecos Texas and only rarely sympatric with other taxa, and the recently described *O. gayleana*, which is found in southeastern New Mexico and adjacent western Texas, with disjunct populations in northern Texas and western Oklahoma (Turner and Moore 2014). Despite low support and low gene tree congruence in our analyses, the two gypsum endemic taxa had moderate sCF support (Fig. 2, S9), much like other taxa with similarly low support and high levels of discordance. In addition,while both gypsum endemics overlapped with other taxa in the morphometric analysis, they occupied morphological extremes in PCA space (Fig. 4). Given that other well-supported taxa also overlap morphologically, it is perhaps not surprising that the two gypsum endemic taxa are not more differentiated from other taxa morphologically. Perhaps the strongest evidence in our data for their recognition as distinct taxa is that we found no evidence of admixture between either of these gypsum endemic taxa and other closely related taxa (Fig. 3a, S10).

### Taxonomic Implications

The most consequential taxonomic result that arises from our analyses is the position of *toumeyi*, a member of subsect. *Salpingia* as circumscribed by Towner (1977). In the present study, this species is resolved as sister to subsect. *Calylophus* with strong support, rendering subsect. *Salpingia* paraphyletic (Fig. 2, S9). Towner (1977) grouped *O. toumeyi* with *O. hartwegii* due to similar floral and bud characters including large flowers and long floral tubes suggestive of hawkmoth pollination, and rounded buds with long, free sepal-tips. Because the breeding system is a defining difference in the current circumscription between the two subsections in sect. *Calylophus*, this result supports abandoning subsections altogether in sect. *Calylophus*.

Within subsect. *Salpingia* our results also suggest the need for revision. While our phylogenetic analyses strongly support the current circumscription of *O. lavandulifolia* (sensu Towner 1977) as a distinct species within subsect. *Salpingia* (Fig. 2, S9), the relationships of the other two species *O. hartwegii* and *O. tubicula* are less clear. Towner (1977) differentiated these two species by the breeding system and grouped the five subspecies of *O. hartwegii* together based on a pattern of reticulate and intergrading variation in which taxa were distinguished from one another by often slight differences in pubescence and leaf shape. Our morphometric analysis confirmed this pattern; however, our phylogenetic results indicated that one taxon, *O. hartwegii* subsp. *fendleri*, shares a closer relationship with the bee pollinated *O. tubicula* subsp. *tubicula* than other taxa in the hawkmoth pollinated *O. hartwegii* species complex (Fig. 2, S6, S7). This relationship was strongly supported and renders *O. hartwegii*, according to the current circumscription, paraphyletic (Towner 1977). Based on strong phylogenetic support for this clade, and its strong morphological distinctiveness as described by Towner (1977), we suggest that *O. hartwegii* subsp. *fendleri* be elevated to the species rank along with both subspecies of *O. tubicula* which were equally well supported in phylogenetic analysis and are geographically isolated. Furthermore, our results support a the possible elevation to species rank for *O. hartwegii* subsp. *filifolia*. While this taxon was poorly supported in our summary coalescent trees (Fig. 2, S6, S7), we found no evidence of hybridization between this taxon and other closely related taxa. In addition, *O. hartwegii* subsp. *fillifolia* is restricted to gypsum. Therefore, we believe that the ecological distinctiveness and lack of gene flow of *O. hartwegii* subsp. *filifolia* with other taxa in the *O. hartwegii* species complex warrants its elevation as a distinct species. In light of these changes, and to maintain consistency in classification in the subsection, we feel that despite the evidence of hybridization of *O. hartwegii* subsp. *pubescens* with *O. hartwegii* subsp. *hartwegii*, it possesses a morphological distinctiveness that is supported by our phylogenetic results. We therefore recommend *O. hartwegii* subsp. *pubescens* be elevated to the species level, while *O. hartwegii* subsp. *hartwegii* and *O. hartwegii* subsp. *maccartii* be retained as is, forming a polytypic species with two subspecies.

In contrast to the relatively clear divisions among taxa in subsect. *Salpingia,* none of the four currently recognized taxa in the subsect. *Calylophus* were consistently recovered as monophyletic. For example, *O. capillifolia* subsp. *capillifolia* was monophyletic in our exon-only summary coalescent tree, but not in the “supercontig” tree, and *O. capillifolia* subsp. *berlandieri* and *O. serrulata* were scattered throughout sect. *Calylophus* in both trees, perhaps suggesting widespread gene flow and/or multiple origins of PTH (Fig. 2, S6). Importantly, our results suggest that the circumscription of *O. gayleana* sensu Turner and Moore (2014) should be amended. Specifically, we find that the populations of subsect. *Calylophus* from northern Texas and western Oklahoma that were assigned to *O. gayleana* by Turner and Moore (2014; *MJM790-5*, *BJC71*) may instead may belong to *O. serrulata* based on both their phylogenetic positions (Fig. 2, S6, S7) and reduced pollen fertility (S12). These north Texas/western Oklahoma populations seem to represent slightly narrower-leaved individuals of *O. serrulata*, which is a common inhabitant of the extensive gypsum outcrops of this area (although it is not restricted to gypsum there).

Finally, our results highlight an unrecognized cryptic taxon within *O. capillifolia* formed by southern Texas coastal populations currently recognized as *O. capillifolia* subsp. *berlandieri*. Towner (1977) described *O. capillifolia* as a polytypic species with two well-differentiated morphological races. Though he noted the geographic and cytological distinction of the southern Texas coastal populations of *O. capillifolia* subsp. *berlandieri*, these populations were included in *O. capillifolia* subsp. *berlandieri* primarily because of completely overlapping morphological variation. In our results, this cryptic clade of southern Texas coastal populations of *O. capillifolia* subsp. *berlandieri* is the most phylogenetically well supported clade in subsect. *Calylophus* and therefore may warrant taxonomic distinction based on our data (Fig. 2, S9r). Similarly, the southern Texas coastal populations of *O. serrulata*, which is likely an independent origin of PTH derived from this cryptic southern Texas coastal clade of *O. capillifolia* subsp. *berlandieri*, are ecologically distinctive and geographically disjunct from other *O. serrulata* (occurring in coastal dunes, unlike other populations in western Texas, Oklahoma, and northern Texas). In the past they were considered distinctive enough to be described as a species, *Calylophus australis* (Towner & Raven 1970). However, Towner (1977) later combined this species with *O. serrulata* based on his decision to treat all PTH populations as *O. serrulata*. Combined with our results here and the ecogeographic distinctiveness consistent with an independent origin of PTH in coastal Texas, we believe this taxon also warrants recognition as a second PTH species in *Oenothera* sect. *Calylophus*.

## CONCLUSIONS

Here we describe a robust example of resolving a recent, rapid radiation using multiple sources of evidence: (1) extensive sampling from populations throughout the geographic and morphological range, (2) target enrichment for hundreds of nuclear genes, (3) the inclusion of flanking non-coding regions, (4) gene tree-based hybridization inference, (5) SNPs extracted from target enrichment data, and (6) morphometrics. Our results indicate that in recently radiated species complexes with low sequence divergence and/or high levels of ILS that could be an intractable problem with traditional loci, the use of targeted enrichment in addition to flanking non-coding regions provides a net benefit and is essential to recover species-level resolution. Our results also underscore the importance of summary coalescent methods and evaluating gene tree discordance for resolving historical relationships in recalcitrant groups. By explicitly testing for hybridization using gene tree approaches, we also demonstrate that the estimated number of character state transitions may be artifactually inflated if hybridization is not taken into account. This, in combination with morphometrics, provided key evolutionary insights where relationships in summary coalescent methods may be obscured by gene flow. Importantly, our study uncovers strong evidence for multiple origins of biologically important phenomena, including the evolution of bee pollination, PTH, and edaphic specialization. Consequently, *Oenothera* sect. *Calylophus* might represent a powerful system for understanding these phenomena, especially with future genome sequence data.

## DATA AVAILABILITY

Illumina reads generated for this study are available at the NCBI Sequence Read Archive under BioProject PRJNA544074. Assembled exon and supercontig sequences, multiple sequence alignments, and SNP files are available at http://dx.doi.org/10.5061/dryad.[NNNN].

## ACKNOWLEDGEMENTS

Material for *O. serrulata* and *O. capillifolia* subsp. *berlandieri* that were used for designing probes for target enrichment (1KP accession codes SJAN and EQTY, respectively) was originally collected by R.A.R.; samples were grown, and RNA was extracted, in the lab of M. Johnson and subsequently sequenced by the One Thousand Plant Transcriptomes (1KP) initiative. We thank the following for access to field sites and permission to collect samples: U.S. Bureau of Land Management (Colorado, New Mexico, Utah), U.S.D.A. Forest Service (Regions 2, 3, and 4), Carlsbad Caverns National Park, Guadalupe Mountains National Park, Big Bend National Park, Palo Duro State Park, and White Sands Missile Range. We thank the Billie L. Turner Plant Resources Center at the University of Texas at Austin for access to herbarium collections, and we thank the following persons for help with collections: David Anderson, George S. Hinton, Nidia Mendoza Díaz, Patrick Alexander, Christopher Martine, Rebecca Drenovsky, Clare Muller, Jeffrey Sanders, Anna Brunner, Joseph Charboneau, Heather-Rose Kates, and Sophia Weinmann. Finally, we thank Elliot Gardner for providing invaluable advice during target enrichment and sequencing, and Daniel Bruzzese for help with collections and for designing the flower silhouette used to signify PTH in figure 1 (available at www.phylopic.org).

## SUPPLEMENTAL MATERIAL

S1.

See excel table “S1 Accessions and Seq Stats”

S2.

Number of leaf tissue accessions sequenced from each taxon

| Taxon | No. of Accessions | 793 |
| --- | --- | --- |
| *O. lavandulifolia* | 16 | 794 |
| *O. tubicula* subsp. *strigulosa* | 3 | 795 |
| *O. tubicula* subsp. *tubicula* | 9 | 796 |
| *O. hartwegii* subsp. *fendleri* | 15 | 797 |
| *O. hartwegii* subsp. *filifolia* | 44 | 798 |
| *O. hartwegii* subsp. *hartwegii* | 12 | 799 |
| *O. hartwegii* subsp. *maccartii* | 9 | 800 |
| *O. hartwegii* subsp. *pubescens* | 26 | 801 |
| *O. hartwegii* | 1 | 802 |
| *O. toumeyi* | 5 | 803 |
| *O. capillifolia* subsp. *berlandieri* | 17 | 804 |
| *O. capillifolia* subsp. *capillifolia* | 11 | 805 |
| *O. gayleana* | 9 | 806 |
| *O. serrulata* | 17 | 807 |
| Outgroups: |  | 808 |
| *O. pilosella* | 1 | 809 |
| *O. organensis* | 1 | 810 |
| *O. primiveris* | 1 | 811 |
| *O. tubifera* | 1 | 812 |
| *O. triloba* | 1 | 813 |
| *O. cespitosa* subsp. *cespitosa* | 1 | 814 |
| *O. suffrutescens* | 1 | 815 |
| *Chylismia scapoidea* subsp. *scapoidea* | 1 | 816 |
| *Eulobus californicus* | 1 | 817 |
| Total | 203 | 818 |

S3.

## MATERIALS AND METHODS

### Taxon Sampling, DNA Extraction, and Determination of PTH

A total of 194 individuals spanning the geographic, morphological, and ecological ranges of all 13 recognized taxa in *Oenothera* sect. *Calylophus* [following Towner (1977) and Turner and Moore (2014)] were included in this study (Fig. 1a, S1, S2) along with eight outgroups representing other major sections of Oenothera (*Eremia*, *Gaura*, *Kneiffia*, *Lavauxia*, *Oenothera*, *Pachylophus*, and *Ravenia*) and other genera (*Chylismia* and *Eulobus*) in Onagraceae (S1, S2). All leaf tissue samples were collected from individuals in the field between 2007 and 2015 and voucher specimens were deposited at the United States National Herbarium (US), with duplicates in most cases at either the Nancy Rich Poole Herbarium (CHIC) or the George T. Jones Herbarium at Oberlin College (OC; S1). DNA was extracted from fresh, silica-dried leaf tissue using either (1) a modified CTAB protocol (Doyle 1987), (2) the Nucleon PhytoPure DNA extraction kit (GE Healthcare Life Sciences, Pittsburgh, Pennsylvania, USA), or (3) a modified CTAB and silicon dioxide purification protocol (Doyle 1987; Sharma and Purohit 2012; See S13 for details) followed by passing any extractions retaining a brown or yellow coloration through a Qiagen Qiaquick PCR spin column for additional purification according to the manufacturer’s protocol (Qiagen, Venlo, Netherlands). The third DNA extraction method was used for difficult to extract, polysaccharide-rich leaf tissue samples that yielded gooey, discolored DNA following initial extraction. PTH status was determined for individuals in subsect. *Calylophus* using floral morphology and/or when flowers were present by assessing pollen fertility using a modified Alexander stain, as PTH taxa have a demonstrated 50% reduction in pollen fertility (Towner 1977). For accessions identified as either *O. capillifolia* subsp. *berlandieri* or *O. serrulata* that had sufficient pollen available, pollen was removed from flowers and stained using a modified Alexander stain (Alexander 1969, 1980). Accessions with less than 50% viable pollen were assigned to *O. serrulata*, the only currently recognized PTH taxon in subsect. *Calylophus*. Pollen count data are provided in Supplement 12 (See S12 for details).

### Bait Design, Library Construction, Target Enrichment, and Sequencing

We targeted 322 orthologous, low-copy nuclear loci determined by clustering transcriptomes of *Oenothera serrulata* (1KP accession *SJAN*) and *Oenothera berlandieri* (1KP accession *EQYT*) to select a subset of the 956 phylogenetically informative *Arabidopsis* loci identified by Duarte et al. (2010). Transcriptomes of two *Oenothera* species, *O. serrulata* and *O. berlandieri*, were assembled and optimal isoforms were filtered for the longest reading frame using custom Perl scripts. Assembled transcripts were aligned as amino acids to the 956 TAIR loci of *Arabidopsis* in TranslatorX (Abascal et al. 2010). This alignment identified 956 orthologous sequences, from which 322 loci were randomly selected. BLAST searches of amino acid sequences from these loci were carried out to ensure orthology between the transcript loci and the *Arabidopsis* TAIR locus. The bait set was designed from these 322 loci, which were selected from both *O. serrulata* and *O. berlanderi* sequences. A set of 19,994 120-bp baits tiled across each locus with a 60 base overlap (2x tiling) was manufactured by Arbor Biosciences (formerly MYcroarray, Ann Arbor, Michigan, USA). Sequencing libraries for 67 samples were prepared with the Illumina TruSeq Nano HT DNA Library Preparation Kit (San Diego, California, USA) following the manufacturer’s protocol, except using half volumes beginning with the second addition of Dynabeads MyOne Streptavidin C1 magnetic beads (Invitrogen, Carlsbad, CA, USA). DNA samples were sheared using a Covaris M220 Focused-Ultrasonicator (Covaris, Woburn, Maryland, USA) to a fragment length of ∼550 bp (for an average insert size of ∼420 bp). The remaining 134 libraries were constructed by Rapid Genomics (Gainesville, Florida, USA), with custom adapters. The Illumina i5 and i7 barcodes were used for all libraries.

Libraries were enriched for these loci using the MyBaits protocol (ArborBiosciences 2016) with combined pools of libraries totaling 1.2 μg of DNA (12 libraries/pool at 100 ng/library). Libraries with less than 100 ng of total recovered DNA were pooled together in equimolar concentrations using available product, resulting in some pools with less than 1.2 μg of DNA. The smallest successful pool contained four samples with 6 ng of library each. Hybridization was performed at 65°C for approximately 18 hours. The enriched libraries were reamplified with 14 to 18 PCR cycles and a final cleanup was performed using a Qiagen QiaQuick PCR cleanup kit following the manufacturer’s protocol to remove bead contamination (Qiagen, Venlo, Netherlands). DNA concentrations were measured using a Qubit 2.0 Fluorometer (Life Technologies, Carlsbad, California, USA) and molarity was measured on an Agilent 2100 Bioanalyzer (Agilent Technologies, Santa Clara, California, USA). A final cleaning step using Dynabeads MyOne Streptavidin C1 magnetic beads was performed on pools with adapter contamination as detected on the Bioanalyzer. Pools were sequenced in four runs at equimolar ratios (4 nM), on an Illumina MiSeq (2 x 300 cycles, v3 chemistry; Illumina, Inc., San Diego, California, USA) at the Pritzker DNA lab (Field Museum, Chicago, IL, USA). This produced 80,273,296 pairs of 300-bp reads. Reads were demultiplexed and adapters trimmed automatically by Illumina Basespace (Illumina 2016). Raw reads have been deposited at the NCBI Sequence Read Archive (BioProject PRJNA544074).

### Quality Filtering, Assembly and Alignment

A summary of read quality from each sample was produced using FastQC (http://www.bioinformatics.babraham.ac.uk/people.html), which revealed read-through adapter contamination in many of the poorer quality samples. To remove read-through contamination and filter for quality, reads were trimmed for known Illumina adapters using Trimmomatic (Bolger et al. 2014) with the following settings: ILLUMINACLIP:<illumina_adapters.fasta>:2:30:10 LEADING:10 TRAILING:10 SLIDINGWINDOW:4:20 MINLEN:20. Trimmed, quality-filtered reads were assembled using the HybPiper pipeline (Johnson et al. 2016) with default settings, followed by the intronerate.py script, to produce both exons and the “splash zone” flanking non-coding region-containing supercontigs. Only pairs with both mates surviving trimming and quality filtering were used for HybPiper.

To compare the influence of “splash-zone” non-coding regions, two sets of alignments were created: (1) exons alone, and (2) coding sequences plus the “splash-zone” (Hereby referred to as supercontigs). For multiple sequence alignments of exons alone, protein and nucleotide sequences assembled in HybPiper were gathered into fasta files by gene. For protein sequences only, stop codons were changed to “X” using a sed command-line regular expression to facilitate alignment, and sequences were aligned using MAFFT with settings: --auto --adjustdirection -- maxiterate 1000 (Katoh et al. 2002). Aligned protein sequences were then used to fit unaligned nucleotide sequences into coding frame alignments using pal2nal with default settings (Suyama et al. 2006). In-frame, aligned DNA sequences were trimmed to remove low-coverage positions and sequences composed only of gaps using TrimAl with the automated setting, which is optimized for maximum likelihood analyses (Gutíerrez et al. 2009). For supercontigs, nucleotide sequences assembled using HybPiper were gathered into fasta files by gene, gene names were removed from fasta headers using a command-line regular expression, and sequences were aligned in MAFFT with settings: --auto --adjustdirection --maxiterate 1000 (Katoh et al. 2002). Reverse compliment tags (“_R_”) were removed from taxon names using a command-line regular expression, and sequences were trimmed using TrimAl with previously listed settings optimized for maximum likelihood analyses (Gutíerrez et al. 2009). To minimize the effects of missing data on phylogenetic analyses, accessions with < 50% of loci passing quality filtering were removed, and genes that were recovered across < 70% of the total remaining samples were also removed. Following quality filtering, we recovered 204 high quality loci (present in at least 70% of samples) and an average of 625,323 reads per sample (S1). Final assemblies are available at Dryad (http://dx.doi.org/10.5061/dryad.[NNNN]). Bioinformatics pipelines and analyses were run at the high-performance computing cluster (“Treubia”) at the Chicago Botanic Garden unless otherwise specified.

### Phylogenetic Reconstruction

We conducted phylogenetic analyses using two strategies for each set of alignments. Alignments were concatenated and analyzed using maximum likelihood (ML) in RAxML (Stamatakis 2014; hereafter referred to as “concatenation”), whereas coalescent-based analyses were conducted using ML gene trees in ASTRAL-II (Mirarab and Warnow 2015; Sayyari and Mirarab 2016) and using unlinked SNPs in SVDquartets (Chifman and Kubatko 2014, 2015) implemented in PAUP* beta version 4.0a168 (Swofford 2003; S3). In concatenation analyses, after aligning each gene separately in MAFFT, genes were concatenated, partitioned, and maximum likelihood trees were reconstructed in RAxML Version 8 (Stamatakis 2014) using the GRTCAT model with 100 “rapid-boostrapping” psuedoreplicates and *Chylismia scapoidea* as the outgroup, on the CIPRES Science Gateway computing cluster (Miller et al. 2010). For coalescent analyses, individual gene trees were first estimated using RAxML Version 8 (Stamatakis 2014), with 100 “rapid-bootstraping” psuedo-replicates and settings: -p 12345 -x 12345 -N 100 -c 25 -f a -m GTRCAT –s, and *Chylismia scapoidea* as the outgroup. Gene trees based on supercontigs were not partitioned by codon position. Coalescent-based analyses of accessions were conducted in ASTRAL-II (Mirarab and Warnow 2015; Sayyari and Mirarab 2016) with default settings using the best RAxML gene trees and their associated bootstrap files as input, and in SVD Quartets (Chifman and Kubatko 2014, 2015) with default settings using an inframe aligned supermatrix with all 204 loci and supercontigs. ASTRAL-II and SVD Quartets analysis was performed with 100 multi-locus bootstraps.

To assess concordance among gene trees and provide additional support complementary to bootstrap values, we conducted two additional analyses. First, we assessed raw gene tree concordance using Phyparts (Smith et al. 2015). Prior to running Phyparts, nodes with < 33% support in the supercontig RAxML gene trees were collapsed using the sumtrees command in Dendropy (Sukumaran 2010). These gene trees were then re-rooted using *Chylismia scapoidea* as the outgroup and ASTRAL-II was rerun using these collapsed, re-rooted gene trees as the input files. Pie charts showing gene tree discordance were generated and overlaid on the resulting ASTRAL-II tree using the PhypartsPiecharts script (https://github.com/mossmatters/phyloscripts/tree/master/phypartspiecharts). Phyparts piecharts and gene tree concordance values were also added to Figure 1 by importing the two data files produced by the Phypartspiecharts.py into R and manually matching them to key nodes on our ASTRAL-II supercontig tree using *ggtree* version 1.14.6 (G Yu, DK Smith, H Zhu, Y Guan 2017). We also generated gene and site concordance factors for our ASTRAL-II tree constructed using supercontigs in IQTree v1.7-beta16 (Minh et al. 2018). IQtree calculates the gene concordance factor (gCF) and accounts for incomplete taxon coverage among gene trees and therefore may provide a more accurate representation of agreement among gene trees than other methods. In addition to gCF, IQTree calculates the site concordance factor (sCF), which is defined as the percentage of decisive nucleotide sites supporting a specific node (Minh et al. 2018). We used the RAxML gene trees produced for the ASTRAL-II supercontig analysis, and supercontig alignments themselves, as the inputs for IQtree. For computing sCF, we randomly sampled 100 quartets around each internal node. Finally, we mapped gCF and sCF values to the ASTRAL-II supercontig tree produced in our previous summary coalescent analysis (Fig. 2). All phylogenetic trees, with the exception of the full Phyparts picharts tree, were visualized using the R package *ggtree* version 1.14.6 (G Yu, DK Smith, H Zhu, Y Guan 2017).

### Ancestral State Reconstruction

To infer ancestral conditions and the number of transitions in reproductive system, we used the *phangorn* (Schliep 2011) package in R. First, a Coalescent-based species tree with accessions grouped into taxa using a mapping file was estimated in ASTRAL-III (Zhang et al. 2018) with default settings using the best RAxML gene trees and their associated bootstrap files, from the supercontig alignments, as input. Next we time calibrated the ASTRAL-III species tree to 1 million years based on estimates of other taxa in the genus (Evans et al. 2009) using the makeChronosCalib function in the *ape* (Paradis et al. 2004) package in R, and estimated an ultrametric tree using the chronos function in *ape* (Paradis et al. 2004) with settings: lambda = 1, model = "relaxed”. Finally, we performed marginal reconstruction of ancestral character states using the maximum likelihood method using the optim.pml and ancestral.pml functions in the *phangorn* (Schliep 2011) package in R.

### Testing for Hybrid Orgins with HyDe

To test for putative hybrid origins of selected taxa, we used HyDe (Blischak et al. 2018) to calculate D-Statistics (Green et al. 2010) for a set of hypotheses (S10). Briefly, HyDe considers a four-taxon network of an outgroup and a triplet of ingroup populations to detect hybridization from phylogenetic invariants that arise under the coalescent model with hybridization. Introgression between P3 and either P1 or P2 influences the relative frequencies of ABBA and BABA, and the D-statistic measures the imbalance between these frequencies. We tested the triplets in (S10) and set *Chylismia scapoidea* as the outgroup. We considered hypothesis tests significant at an overall α < 0.05 level with estimates of γ between 0 and 1. Z-scores greater than 3 are generally interpreted as strong evidence of introgression.

### Population-level Analysis

To further characterize population-level processes or genetic structure within sect. *Calylophus*, we extracted and filtered SNPs by mapping individual reads against reference supercontigs (see https://github.com/lindsawi/HybSeq-SNP-Extraction). To account for duplicates arising from PCR during HybSeq in SNP calling and filtering, first we selected the sample with the highest target recovery rate and sequencing depth as a target reference sequence (Oenothera_capillifolia_berlandieri_bjc19) and gathered supercontigs for this individual into a single target FASTA file. We then ran BWA (Li and Durbin 2009) to align sequences, Samtools ‘index’ (Danecek et al. 2011) and GATK CreateSequenceDictionary (Poplin et al. 2017), respectively, on the resulting target FASTA file. Next we ran a custom script “variant_workflow.sh” using both read files from each *Calylophus* sample as input to create a vcf file for each sample. SNP’s were called for each individual using GATK (Poplin et al. 2017) and the vcf file from each sample as input. The resulting vcf file created in the previous step was filtered to remove indels using GATK and the original target FASTA file as input, and then filtered again based on read mapping and quality with GATK VariantFiltration with settings: -- filterExpression "QD < 5.0 || FS > 60.0 || MQ < 40.0 || MQRankSum < -12.5 || ReadPosRankSum < -8.0" (Poplin et al. 2017). Finally, we generated a reduced SNP file in FastStructure format using PLINK (Purcell et al. 2007) to remove SNPs that did pass filter using the command: plink - -vcf-filter --vcf Pachylophus.filtered.snps.vcf --const-fid --allow-extra-chr --geno --make-bed -- recode structure. Finally we used Discriminant Analysis of Principal Components (Jombart et al. 2010) as implemented in the R package *adegenet* (Jombart 2008) and the snmf function in the LEA package (Frichot and François 2015) in R (R Core Team, 2020).

### Morphological Measurements and Analysis

To assess taxon boundaries and patterns of morphological variation, we measured character states for the following key morphological structures that have been used historically to discriminate taxa in sect. *Calylophus* (Towner 1977): plant height, leaf length (distal), leaf width (distal), leaf length/width ratio (distal), leaf length (basal), leaf width (basal), leaf length/width (basal), sepal length, and sepal tip length. Measurements were made with digital calipers when possible, or with a standard metric ruler and dissecting scope, from voucher specimens of nearly all sampled populations of sect. *Calylophus* included in our molecular phylogenetic analyses. Measurements are provided in S11 and have been deposited at Dryad (http://dx.doi.org/10.5061/dryad.[NNNN]). Morphological measurements were log transformed using the R base function ‘log’ (R Core Team 2018) prior to Principal Components Analysis (PCA), which was conducted in R using the *stats* package version 3.7 and the function ‘prcomp’(R Core Team 2018). All ‘NA’ values were omitted from analysis. Plots of PCA results were visualized using the *ggplot2* package in R (Wickham 2016).

---

S4.

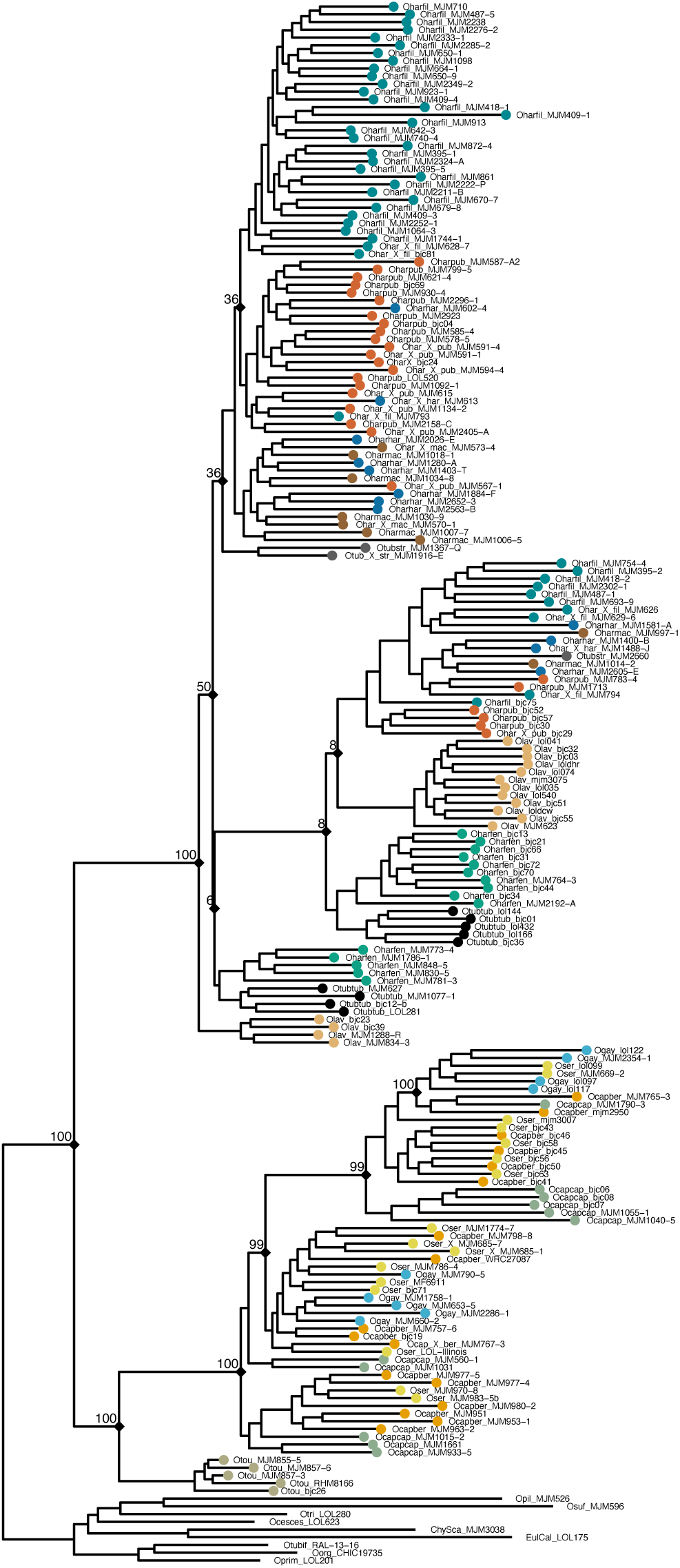
Concatenation tree constructed using the exon-only dataset and 100 bootstraps. Bootstrap values indicated at relevant nodes.

S5.

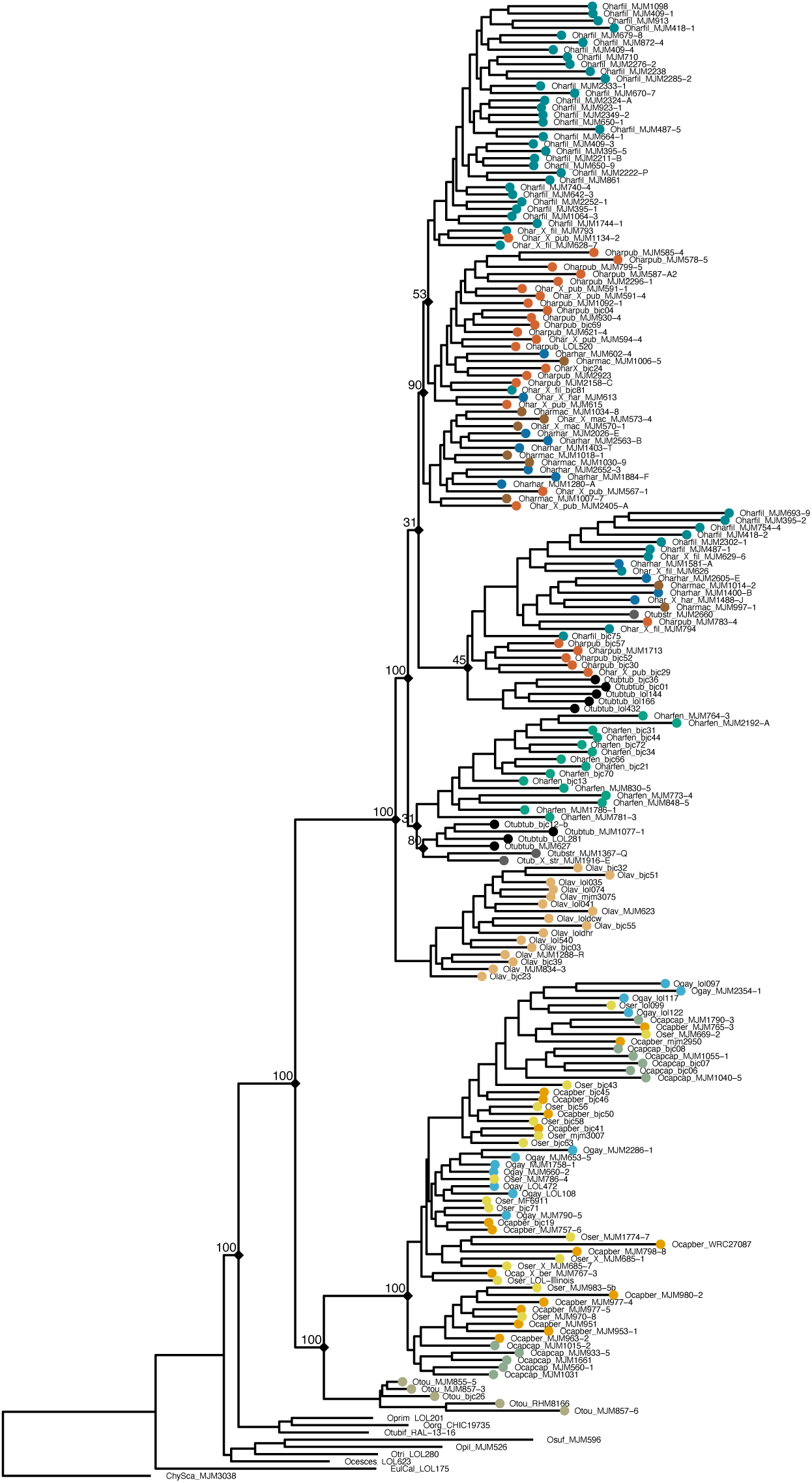
Concatenation tree constructed using the supercontig dataset and 100 bootstraps. Bootstrap values indicated at relevant nodes.

S6.

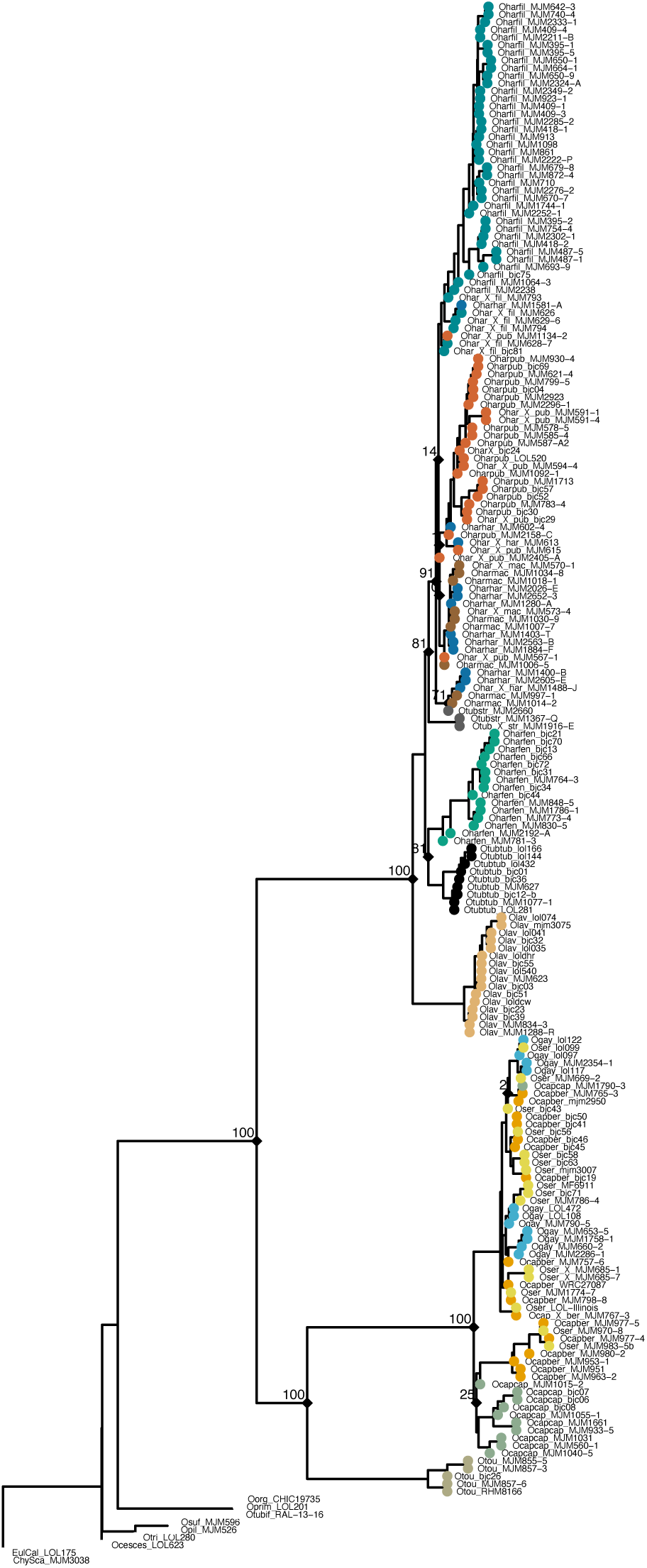
ASTRAL-II summary coalescent tree constructed using the exon-only dataset and 100 bootstraps. Bootstrap values indicated at relevant nodes.

S7.

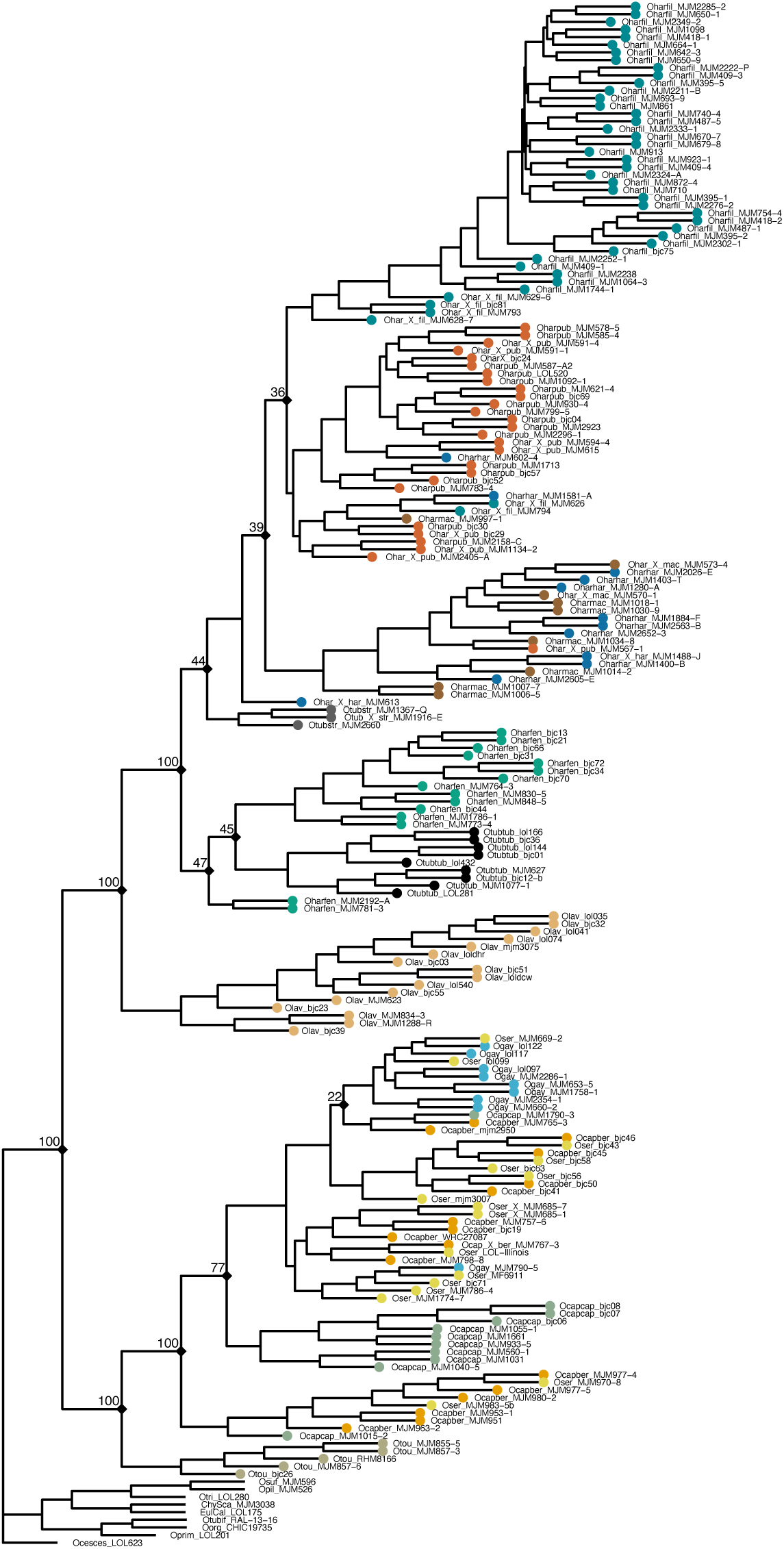
SVD Quartets summary coalescent tree constructed using the supercontig dataset with 100 bootstraps. Bootstrap values indicated at relevant nodes.

S8.

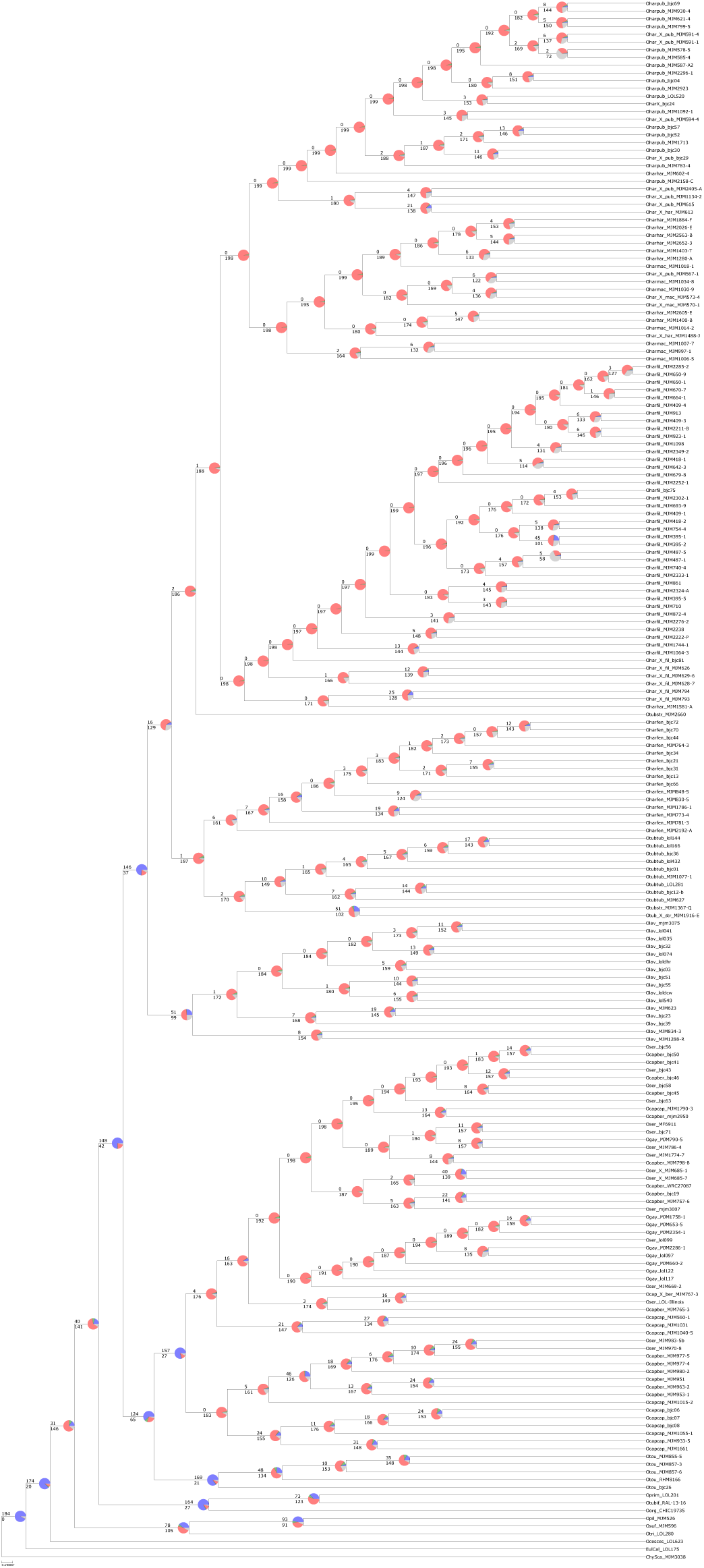
Phyparts piecharts ASTRAL-II tree constructed using the supercontig dataset. Piechart colors correspond to: blue = concordant, green = top alternative bipartition, red = all other alternative bipartitions, black = uninformative for that node.

S9.

Summary of support values for current and proposed taxonomic treatments, by analysis. ‘e’ signifies trees based on exon-only data, “e+i” signifies trees based on supercontigs. ‘*p’* indicates paraphyletic, unsupported taxon treatment according to tree topology.

| Taxon | Concat. e | Concat. e+i | ASTRAL-II e | ASTRAL-II e+i | SVDQ e+i | Phyparts | gCF iQtree e+i | sCF iQtree e+i |
| --- | --- | --- | --- | --- | --- | --- | --- | --- |
| a. *Oenothera hartwegii* (Towner 1977) | *p* | *p* | *p* | *p* | *p* | *p* | *p* | *p* |
| b. *Oenothera hartwegii* subsp. *fendleri* (Towner 1977) | <50 | *p* | 81 | 99 | *p* | 1/ 188 | 1 | 38 |
| c. *Oenothera hartwegii* subsp. *filifolia* (Towner 1977) | *p* | *p* | <50 | <50 | <50 | 1/ 190 | 0 | 36 |
| d. *Oenothera hartwegii* subsp. *hartwegii* (Towner 1977) | *p* | *p* | *p* | *p* | *p* | *p* | *p* | *p* |
| e. *Oenothera hartwegii* subsp. *macartii* (Towner 1977) | *p* | *p* | *p* | *p* | *p* | *p* | *p* |  |
| f. *Oenothera hartwegii* subsp. *hartwegii* + subsp. *macartii* | *p* | *p* | <50 | <50 | <50 | 0/ 200 | 0 | 33 |
| g. *Oenothera hartwegii* subsp. *pubescens* (Towner 1977) | *p* | *p* | <50 | <50 | <50 | 0/ 200 | 0 | 36 |
| h. *Oenothera lavandulifolia* (Towner 1977) | 100 | *p* | 100 | 100 | 100 | 147/37 | 74 | 83 |
| i. *Oenothera tubicula* (Towner 1977) | *p* | *p* | *p* | *p* | *p* | 1/ 188 | *p* | *p* |
| j. *Oenothera tubicula* subsp. *tubicula* (Towner 1977) | *p* | *p* | 81 | 99 | <50 | 2/ 171 | 1 | 38 |
| k. *Oenothera tubicula* subsp. *strigulosa* (Towner 1977) | *p* | *p* | 81 | 100 | <50 | *p* | 0 | 37 |
| l. *Oenothera fendleri* + *Oenothera tubicula* subsp. *tubicula* | *p* | *p* | 100 | 100 | <50 | 16/129 | 10 | 40 |
| m. *Oenothera toumeyi* (Towner 1977) | 100 | 100 | 100 | 100 | 100 | 125/65 | 62 | 56 |
| n. *Oenothera toumeyi* + subsect. *Calylophus* | 100 | 100 | 100 | 100 | 100 | 149/42 | 76 | 65 |
| o. *Oenothera capillifolia* (Towner 1977) | *p* | *p* | *p* | *p* | *p* | *p* | *p* | *p* |
| p. *Oenothera capillifolia* subsp. *capillifolia* (Towner 1977) | *p* | *p* | <50 | *p* | <77 | *p* | *p* | *p* |
| q. *Oenothera capillifolia* subsp. *berlandieri* (Towner 1977) | *p* | *p* | *p* | *p* | *p* | *p* | *p* | *p* |
| r. *Oenothera capillifolia* subsp. *berlandieri* (New Taxon; South Texas coastal populations) | *p* | *p* | <50 | 100 | 100 | 157/27 | 79 | 86 |
| s. *Oenothera serrulata* (Towner 1977) | *p* | *p* | *p* | *p* | *p* | *p* | *p* | *p* |
| t. *Oenothera* ‘*australis*’ (New Taxon; South Texas *O. serrulata*) | *p* | 100 | *p* | *p* | *p* | 10/175 | *p* | *p* |
| u. *Oenothera gayleana* (Turner & Moore 2014) | *p* | *p* | *p* | *p* | *p* | *p* | *p* | *p* |
| v. *Oenothera gayleana* (Revised Taxon) | *p* | *p* | *p* | <50 | <50 | 0/ 193 | 0 | 35 |

S10.

List of admixture hypotheses tested, Zscore, P-value and Gamma results from HyDe analysis. Each row represents a triplet set that was tested consisting of a putative hybrid individual and two parent groups; Parent 1 and Parent 2 (see “HyDe Group” Appendix I for group membership).

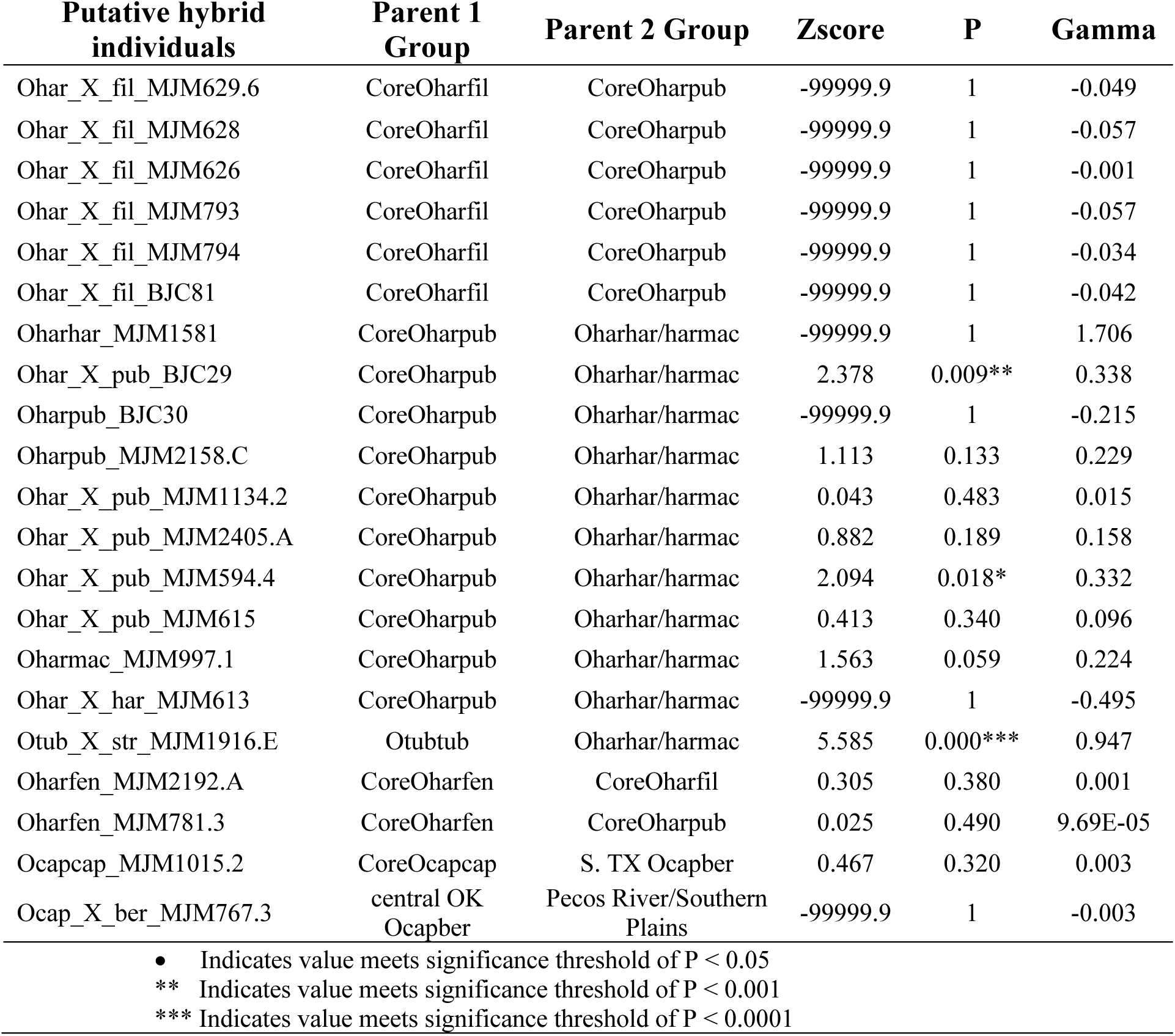

S11.

See excel table “S11 Morphometric Data”.

S12.

See excel table “S12 Pollen Counts”.

S13.

## Oenothera HybSeq CTAB Silica DNA extractions b. cooper 11/10/2015

Adapted from JJSA Protocol, J. Fant’s Protocol and Sharma and Puohit (2012) for Silica-dried Leaves of *Oenothera* sect. *Calylophus*

### SAMPLE PREPARATION & GRINDING

1. Warm <u>CTAB buffer</u> to 65°C in water bath in the hood and pour liquid nitrogen into dipping canister (You will only need a few inches of liquid in the Dewar, make sure to pour it back into the large holding canister when you are done to prevent evaporation).
2. Put a small volume of 1:1 sterilized sand and 3 metal/ceramic beads into centrifuge tubes, then put tubes in freezer (-20) until envelopes are ready.
3. Keeping envelopes/leaf tissue on ice to prevent thawing; take envelopes and centrifuge tubes out of freezer and weigh/estimate small amount of dried tissue (e.g., 10-50 mg of leaves), put in labeled microfuge tubes. After each tube is filled, place tube in freezer.

a. Make sure leaf tissue is broken up into small pieces when adding to ensure full grinding.
b. Keeping tissue frozen prevents phenolic compounds from oxidizing, and enzymes from activating and breaking down DNA
c. Follow DNA free lab protocols using bleach to prevent contamination/cross contamination of samples.
4. Mix 45 (= .3%) µl 2-mercaptoethanol to 15 ml <u>CTAB buffer</u> in small beaker (enough for ∼15 samples…double for 30 samples, etc.)

a. Mercaptoethanol must be measured and added in the fume hood.
b. CTAB is a detergent that lyses (breaks down) the cell membranes. The PVP in the buffer helps bind the polysaccharides and might co-ppt
5. Dip tubes in Liquid Nitrogen (or store in -80 for ∼30 minutes) and **immediately** Grind tissue **<u>thoroughly</u>** in Fast Prep machine (thorough grinding may require 2-4 cycles, re-dip tubes in liquid nitrogen after 2 cycles to ensure no thawing occurs).
6. **Immediately** after grinding add 600 µl of CTAB/mercaptoethanol buffer to each tube

a. **BE CAREFUL.** It is important to prevent cross-contamination when opening tubes. **Powdered tissue can spread easily through the air or on your fingers.** Make sure open tubes are spaced apart from one another on the tray; open each tube away from other open tubes/caps; and place caps a safe distance from one another. Make sure leaf powder is not stuck to your gloves before touching consecutive tubes/caps.
b. Grind on FastPrep for one cycle

### EXTRACTION AND ISOLATION OF DNA (once leaf tissue is well-ground)

1. Incubate tubes for 60 minutes at 60°C on Thermoblock or water bath

a. Shake tubes several times ∼ every 10 minutes during incubation (for compacted samples, vortex on high setting to ensure mixing)
b. When incubation is complete, turn Thermoblock down to 37°C
2. Add 600 µL chloroform-isoamyl (24:1) to tube and vortex on High setting to ensure thorough mixing.

a. **BE CAREFUL.** It is important to prevent cross-contamination at this step when opening/handling tubes. Follow steps outlined previously.
b. Chlorophyll and other pigments get transferred into the CI layer because pigments are non-polar and dissolve into the highly non-polar chloroform. The DNA remains in the aqueous layer.
3. Spin for 10 min at 9,000 RPM.

a. After spinning, the upper layer of each tube should have a clear liquid: this is where your DNA is.
4. Transfer top (clear) liquid to a new (labeled) 1.5 ml eppindorf tube. **Do not be Greedy**, leaving some liquid in tube is fine – the important thing is not to pipette **ANY** interphase material.

a. A yield of ∼400-500 µL of supernatant is common.
b. If you accidently disturb the interphase, or believe that you have pipetted interphase, return supernatant to microfuge tube and re-centrifuge for 10 min at 9000 rpm. Repeat step 4.

### DNA PRECIPITATION

1. Add 1/10 volume 3M sodium acetate and ½ volume 5M NaCl (i.e. 500ul of solution add 250 ul of NaCl, 50 ul of Na-Acetate).
2. Add 2/3^rd^ volume of cold isopropanol (to previous volumes = 500 ul isop)

a. Isoproponal is heavy alcohol so allows DNA ppt with lower volume
3. Mix & store in -20°C (freezer) for 1-2 hours

***(NOTE: POTENTIAL STOPPING POINT FOR A DAY OR MORE…SAMPLES CAN STAY AT -20°C FOR DAYS)***

4. Spin for 3 min at **10K-12K RPM.**
5. Decant the supernatant & drain; make sure pellet stays at bottom of tube.

a. Pellets are usually white but sometimes they can even be brown, this is not necessarily a problem.
6. Wash pellet with 500ul of <u>70</u>% EtOH & flick tubes to dislodge pellet from bottom, then vortex on high to clean.
7. Spin for 3 min at 10K-12K RPM.
8. Decant the supernatant making sure pellet stays at bottom of tube
9. Leave tubes open and place tubes on Speedvac (vacuum on -2.5, heat = low, spin) for ∼20 minutes or until all EtOH has evaporated.

a. If pellets are not dry after 20 minutes, check frequently to prevent over drying which makes the pellet difficult to re-suspend.
b. Alternatively, can air dry on thermoblock

***(NOTE: POTENTIAL STOPPING POINT …SAMPLES CAN STAY AT 20°C FOR DAYS – place in cupboard but cover with kimwipe)***

### FINAL CLEANING & RESUSPENSION OF DNA

1. Add 500 ul of 0.5 mL High Salt TE Buffer and close lids
2. Incubate tubes on Thermoblock at 37-50°C (45) until pellet dissolves (15-30 minutes); Vortex frequently if this is taking a long time

***(NOTE: POTENTIAL STOPPING POINT FOR A DAY OR MORE…SAMPLES CAN STAY AT -20°C FOR DAYS)***

3. Add 3 volumes of binding buffer (3 volumes for every 1 volume DNA pellet = ∼100-150 ul BB), let stand for 20 minutes
4. Centrifuge at 550 g for 10 min
5. transfer supernatant to clean 2ml eppi tube leaving any colored or gelatinous precipitate behind.
6. Add 300 µl of silica suspension and mix for 30 minutes by regular and frequent gentle inversion
7. Centrifuge at 550 g for 10 minutes, discard supernatent
8. Re-suspend silica pellet in 1.5 mL of wash buffer 1
9. Centrifuge at 3000 g for 15 seconds, decant supernatent
10. Re-suspend silica pellet in 1.5 mL of wash buffer 2
11. Centrifuge at 3000 g for 15 seconds, decant supernatant, dry pellet completely on speed vac
12. suspend silica pellet in 300 ul TE buffer, incubate at 50° mixing regularly by vortex
13. centrifuge at 11600 g for 1 min. then transfer supernatant into clean 1.5 mL eppi tube.
14. Precipitate DNA by adding 50 ul of 3M sodium Acetate and 500 uL 100% Etoh
15. Store at -20° C for at least 2 hours

***(NOTE: POTENTIAL STOPPING POINT FOR A DAY OR MORE…SAMPLES CAN STAY AT -20°C FOR DAYS)***

16. Centrifuge at 11600 g for 5 min, discard supernatant, dry pellet on speed vac
17. Dissolve in 50 uL 1X TE buffer (it may not look like there is anything in the tube at this point, but there is lots of pure DNA in there! Add the TE buffer)
18. Store in -20°C (freezer)
19. In rare circumstances that an extraction still has coloration at this stage (yellow or brown usually) you can now put it through a Qiagen Qiaquick spin column or other proprietary cleanup column to further purify the DNA.

### REAGENTS (recipes below) and SUPPLIES

• CTAB buffer (in glass container in refrigerator)
• Liquid Nitrogen
• 2-mercaptoethanol (in fume hood with gloves)
• 24:1Chloroform:isoamyl alcohol - glass container in fumehood use gloves and keep in hood -(Or 25:24:1 phenol-chloroform:isoamyl)
• 5M Nacl
• 3M M sodium acetate
• Isopropanol and 70% and 100% EtOH
• High Salt TE buffer
• Binding Buffer
• Silica Matrix
• Wash Buffer 1
• Wash Buffer 2
• 1X TE buffer
• 1.5 ml eppi tubes
• 2 ml eppi tubes

**CTAB BUFFER** [100 mM Tris-HCl (pH=8.0), 1.4 M NaCl, 20 mM EDTA, 2% (or 4%) CTAB

(hexadecyltrimethylammonium bromide), 2% PVP-40 (polyvinylpyrollidone, m.w. 40000), + 0.3% β- mercaptoethanol (add in fume hood in a beaker to a premeasured volume of CTAB buffer required for the number of extractions planned plus one).]

For 200 mL CTAB buffer 2% (4%):

20 ml 1M Tris-Cl

1.4 M NaCl (16.36 g NaCl)

• The addition of NaCl at concentrations higher than 0.5 M, along with CTAB, is known to remove polysaccharides during DNA extraction

8 ml 0.5 M EDTA

4 g CTAB (8 g for 4%)

4 g PVP-40

∼120 ml dH2O, then fill to 200 mL

**TE buffer**: [10 mM Tris-HCl (pH=8.0), 1 mM EDTA.]

10 mL 1 M Tris-HCl

2 ml 0.5 M EDTA

982 ml dH2O

**High Salt TE Buffer**: [10 mM Tris-HCl (pH=8.0), 1 mM EDTA, 1M NaCl.]

10 ml 1 M Tris-HCl

2 ml 0.5 M EDTA

200 ml 5M NaCl

788 ml dH2O

**Binding Buffer**: [50mM Tris (pH7.5), 6M NaClO4, 1mM EDTA]

For 200 ml:

10mL Tris

168.552 g NaClO4

400 ul EDTA

top off to 200 mL with dH20

**Wash Buffer 1**: [3 volumes binding buffer, 1 volume water]

For 60 samples:

90 ml Binding Buffer

30 ml dH20

**Wash Buffer 2**: [1 volume 40 mM Tris (pH 8.0), 4 mM EDTA, 0.8 M NaCl, 1 Volume ethanol]

For 60 samples:

1.8 ml 1M tris

360 ul .5M EDTA

7.2 ml 5M NaCL

35.64 ml dh20

35.65 45 ml 100% etoh

**24:1 Chloroform: Isoamyl**

240ml Chloroform

10ml Isoamyl alcohol

25:24:1 Chloroform: Isoamyl Saturated with 10 mM Tris, pH 8.0, 1 mM EDTA. – Purchase to ensure right pH

250ml Phenol

240ml Chloroform

10ml Isoamyl alcohol

**Silica suspension (use lab grade silicon dioxide 99%)**

1. Suspend silicon dioxide powder in ∼20 volumes of water (50 ml : 1L)

a. use magnetic stirring rod to bring silica into uniform suspension
2. Decant suspension into clean beaker leaving behind heavy sediment at bottom (discard this)
3. Allow suspension to settle by gravity for 2-3 hours
4. Decant and discard supernatant with fine material keeping silicon dioxide that has settled
5. Re-suspend silicon dioxide in dH20, repeating step until fine particles removed (2-3X total is good)

a. use magnetic stirring rod to bring silica into uniform suspension
6. Re-suspend silicon dioxide in dH20 and transfer to 15 ml falcon tubes
7. Collect silicon dioxide by low speed centrifugation (up to 3200 rpm for 3 min)
8. Add ∼15 ml of 10% bleach solution to each aliquot and mix frequently by inversion/vortexing for 15-30 minutes
9. Re-suspend diatomite in dH20 by vortexing, then centrifuge at low speed for 3 minutes to pellet
10. Discard supernatent
11. Repeat step 9 and 10 two more times
12. Re-suspend in dH20 and autoclave
13. Re-suspend silica dioxide pellet in 1 volume PCR grade DNA/RNA free H2O
14. Aliquot into 1.5 ml eppi tubes and store at 4° C
15. Vortex thoroughly to bring silica into uniform suspension before using

---

s14.

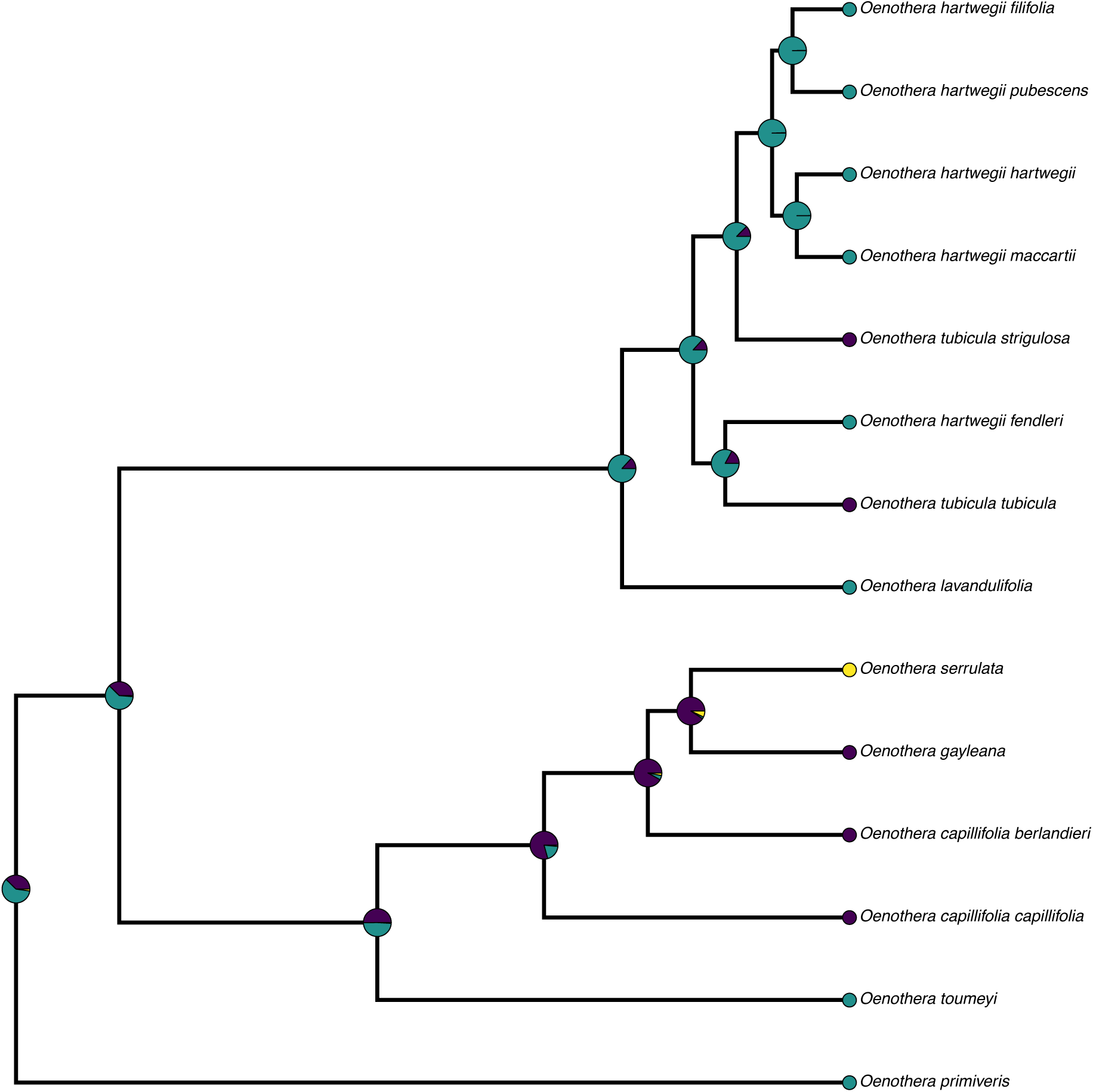
Ancestral State Reconstruction of reproductive system in sect. *Calylophus* using supercontigs and accessions grouped into taxa. Pie-charts on nodes represent likelihood of ancestral reproductive system at each node (teal = hawkmoth pollination, purple = bee pollination, yellow = PTH).

## REFERENCES

Alexander M.P. 1969. Differential Staining of Aborted and Nonaborted Pollen. Stain Technol. 44:117–122.

Alexander M.P. 1980. A Versatile Stain for Pollen Fungi, Yeast and Bacteria. Stain Technol. 55:13–18.

Anacker B.L., Whittall J.B., Goldberg E.E., Harrison S.P. 2011. Origins and consequences of serpentine endemism in the California flora. Evolution (N. Y). 65:365–376.

Artz D.R., Villagra C. A., Raguso R. A. 2010. Spatiotemporal variation in the reproductive ecology of two parapatric subspecies of *Oenothera cespitosa* (Onagraceae). Am. J. Bot. 97:1498–510.

Barrett S., Harder L., Worley A. 1996. The comparative biology of pollination and mating in floweing plants. Philos. Trans. R. Soc. B Biol. Sci. 351:1271–1280.

Barrett S.C.H. 2013. The evolution of plant reproductive systems: how often are transitions irreversible? Proc R Soc B. 280:20130913.

Barthell J. F., Knops J. M. 1997. Visitation of evening primrose by carpenter bees: evidence of a" mixed" pollination syndrome. Southwest. Nat. 86–93.

Blischak P.D., Chifman J., Wolfe A.D., Kubatko L.S. 2018. HyDe: A python package for genome-scale hybridization detection. Syst. Biol. 67:821–829.

Boberg E., Alexandersson R., Jonsson M., Maad J., Ågren J., Nilsson L.A. 2014. Pollinator shifts and the evolution of spur length in the moth-pollinated orchid *Platanthera bifolia*. Ann. Bot. 113:267–275.

Bolger A.M., Lohse M., Usadel B. 2014. Trimmomatic: A flexible trimmer for Illumina sequence data. Bioinformatics. 30:2114–2120.

Brady K.U., Kruckeberg A.R., Bradshaw H.D. 2005. Evolutionary ecology of plant adaptation to serpentine soils. Annu. Rev. Ecol. Evol. Syst. 36:243–266.

Bruzzese D.J., Wagner D.L., Harrison T., Jogesh T., Overson R.P., Wickett N.J., Raguso R.A., Skogen K.A. 2019. Phylogeny, host use, and diversification in the moth family Momphidae (Lepidoptera: Gelechioidea). PLoS One. 14:e0207833.

Bryson R.W., Linkem C.W., Dorcas M.E., Lathrop A., Jones J.M., Alvarado-Díaz J., Grünwald C.I., Murphy R.W. 2014. Multilocus species delimitation in the *Crotalus triseriatus* species group (serpentes: Viperidae: Crotalinae), with the description of two new species. Zootaxa. 3826:475–496.

Cacho N.I., Strauss S.Y. 2014. Occupation of bare habitats, an evolutionary precursor to soil specialization in plants. Proc. Natl. Acad. Sci. 111:15132–15137.

Campbell D.R.., Waser N.M.., Melendez-Ackerman E.J.. 1997. Analyzing pollinator-mediated selection in a plant hybrid zone: Hummingbird visitation patterns on three spatial scales. Am. Nat. 149:295–315.

Chifman J., Kubatko L. 2014. Quartet inference from SNP data under the coalescent model. Bioinformatics. 30:3317–3324.

Chifman J., Kubatko L. 2015. Identifiability of the unrooted species tree topology under the coalescent model with time-reversible substitution processes, site-specific rate variation, and invariable sites. J. Theor. Biol. 374:35–47.

Christie M.R., Knowles L.L. 2015. Habitat corridors facilitate genetic resilience irrespective of species dispersal abilities or population sizes. Evol. Appl. 8:454–463.

Crepet W.L., Niklas K.J. 2009. Darwin’s second “abominable mystery”: Why are there so many angiosperm species? Am. J. Bot. 96:366–81.

Duarte J.M., Wall P.K., Edger P.P., Landherr L.L., Ma H., Pires J.C., Leebens-Mack J., dePamphilis C.W. 2010. Identification of shared single copy nuclear genes in *Arabidopsis, Populus, Vitis* and *Oryza* and their phylogenetic utility across various taxonomic levels. BMC Evol Biol. 10:61.

Eckert A.J., Carstens B.C. 2008. Does gene flow destroy phylogenetic signal? The performance of three methods for estimating species phylogenies in the presence of gene flow. Mol. Phylogenet. Evol. 49:832–842.

Ehrlich, P. R., & Raven, P. H. (1969). Differentiation of populations. Science. 165:1228–1232. Folk R.A., Mandel J.R.,

Freudenstein J. V. 2015. A protocol for targeted enrichment of intron-containing sequence markers for recent radiations: A phylogenomic example from Heuchera (Saxifragaceae). Appl. Plant Sci. 3:1500039.

Frichot E, Mathieu F, Trouillon T, Bouchard G and François O (2014) Fast and efficient estimation of individual ancestry coefficients. Genetics. 196:973–983.

Gerard D., Gibbs H.L., Kubatko L. 2011. Estimating hybridization in the presence of coalescence using phylogenetic intraspecific sampling. BMC Evol. Biol. 11:291.

Giarla T.C., Esselstyn J.A. 2015. The challenges of resolving a rapid, recent radiation: Empirical and simulated phylogenomic of Philippine shrews. Syst. Biol. 64(5):727–740.

Green R., Krause J., Briggs A., Rasilla Vives M., Fortea Pérez F. 2010. A draft sequence of the neandertal genome. Science. 328:710–722.

Heyduk K., Trapnell D.W., Barrett C.F., Leebens-mack J.I.M. 2016. Phylogenomic analyses of species relationships in the genus *Sabal* ( Arecaceae ) using targeted sequence capture. Biol. J. Linn. Soc. 117(1):106–120.

Hollister J.D., Greiner S., Johnson M.T.J., Wright S.I. 2019. Hybridization and a loss of sex shape genome-wide diversity and the origin of species in the evening primroses (Oenothera, Onagraceae). New Phytol. 224:1372–1380.

Jogesh T., Overson R.P., Raguso R.A., Skogen K.A. 2017. Herbivory as an important selective force in the evolution of floral traits and pollinator shifts. AoB Plants. 9(1):plw088.

Johnson M.G., Gardner E.M., Liu Y., Medina R., Goffinet B., Shaw A.J., Zerega N.J.C., Wickett N.J. 2016. HybPiper: Extracting coding sequence and introns for phylogenetics from high-throughput sequencing reads using target enrichment. Appl. Plant Sci. 4:1600016.

Johnson M.T.J., Smith S.D., Rausher M.D. 2009. Plant sex and the evolution of plant defenses against herbivores. Proc. Natl. Acad. Sci. 106:18079–18084.

Jombart, T. (2008) adegenet: a R package for the multivariate analysis of genetic markers. Bioinformatics 24:1403–1405.

Jombart T, Devillard S and Balloux, F (2010). Discriminant analysis of principal components: a new method for the analysis of genetically structured populations. BMC Genetics. 11:94.

Katinas L., Crisci J., Wagner W., Hoch P. 2004. Geographical diversification of tribes Epilobieae, Gongylocarpeae, and Onagreae (Onagraceae) in North America, based on parsimony analysis of endemicity and track compatibility analysis. Ann. Missouri Bot. 91:159–185.

Knowles L.L. 2009. Estimating species trees: Methods of phylogenetic analysis when there is incongruence across genes. Syst. Biol. 58:463–467.

Knowles L.L., Chan Y.-H. 2008. Resolving species phylogenies of recent evolutionary radiations. Ann. Missouri Bot. Gard. 95:224–231.

Kruckeberg A. 1984. California Serpentines: Flora, Vegetation, Geology, Soils, and Management Problems. Berkeley: Univ of California Press.

Kubatko L.S., Chifman J. 2019. An invariants-based method for efficient identification of hybrid species from large-scale genomic data. BMC Evol. Biol. 19:1–13.

Leaché A.D., Harris R.B., Rannala B., Yang Z. 2014. The influence of gene flow on species tree estimation: a simulation study. Syst. Biol. 63:17–30.

Lemmon A.R., Emme S.A., Lemmon E.M. 2012. Anchored hybrid enrichment for massively high-throughput phylogenomics. Syst. Biol. 61:727–744.

Levin R., Wagner W., Hoch P. 2004. Paraphyly in Tribe Onagreae : Insights into Phylogenetic Relationships of Onagraceae Based on Nuclear and Chloroplast Sequence Data. Syst. Bot. 29:147–164.

Lewis, Emily. 2015. Differences in Population Genetic Structure of Hawkmoth and Bee-Pollinated Species of Oenothera (Onagraceae) Are More Pronounced at a Landscape Scale. Northwestern University Libraries. Masters Thesis. Northwestern University.

Maddison W.P., Knowles L. 2006. Inferring Phylogeny Despite Incomplete Lineage Sorting. Evol. Biol. 55:21–30.

Mamanova L., Coffey A.J., Scott C.E., Kozarewa I., Turner E.H., Kumar A., Howard E., Shendure J., Turner D.J. 2010. Target-enrichment strategies for next-generation sequencing. Nat. Methods. 7:111–118.

Mandel J.R., Dikow R.B., Funk V. A, Masalia R.R., Staton S.E., Kozik A., Michelmore R.W., Rieseberg L.H., Burke J.M. 2014. A target enrichment method for gathering phylogenetic information from hundreds of loci: An example from the Compositae. Appl. Plant Sci. 2:1– 6.

Meng C., Kubatko L.S. 2009. Detecting hybrid speciation in the presence of incomplete lineage sorting using gene tree incongruence: A model. Theor. Popul. Biol. 75:35–45.

Miller R.B.. 1981. Hawkmoths and the Geographic Patterns of Floral Variation in *Aquilegia caerulea*. Evolution (N. Y). 35:763–774.

Minh B.Q., Hahn M.W., Lanfear R. 2020. New methods to calculate concordance factors for phylogenomic datasets. Mol. Biol. Evol. 37(9):2727–2733.

Mirarab S., Warnow T. 2015. ASTRAL-II: Coalescent-based species tree estimation with many hundreds of taxa and thousands of genes. Bioinformatics. 31:i44–i52.

Miyake T., Yahara T. 1998. Why does the flower of *Lonicera japonica* open at dusk?. Can. J. Bot. 76(10):1806–1811.

Moore M., Jansen R. 2007. Origins and biogeography of gypsophily in the Chihuahuan Desert plant group *Tiquilia* Subg. subg. *Eddya*. Syst. Bot. 32:392–414.

Moore M.J., Mota J.F., Douglas N.A., Flores Olvera H., Ochoterena H. 2014. Ecology assembly evolution gypsophile floras. In: Rajakaruna N., Boyd R.S., Harris T.B., editors. Plant Ecology and Evolution in Harsh Environments. New York: Nova Science Publishers, Inc. p. 97–128.

Nason J.D., Hamrick J.L., Fleming T.H. 2002. Historical vicariance and postglacial colonization effects on the evolution of genetic structure in *Lophocereus*, a Sonoran Desert columnar cactus. Evolution 56:2214–2226.

Van der Niet T., Johnson S.D. 2012. Phylogenetic evidence for pollinator-driven diversification of angiosperms. Trends Ecol. Evol. 27:353–361.

van der Niet T., Johnson S.D., Linder H.P. 2006. Macroevolutionary data suggest a role for reinforcement in pollination system shifts. Evolution 60:1596–1601.

Van Der Niet T., Peakall R., Johnson S.D. 2014. Pollinator-driven ecological speciation in plants: New evidence and future perspectives. Ann. Bot. 113:199–211.

Peakall R., Ebert D., Poldy J., Barrow R.A., Francke W., Bower C.C., Schiestl F.P. 2010. Pollinator specificity, floral odour chemistry and the phylogeny of Australian sexually deceptive *Chiloglottis* orchids: implications for pollinator-driven speciation. New Phytol. 188:437–50.

Rajakaruna N. 2004. The Edaphic Factor in the Origin of Plant Species. Int. Geol. Rev. 46:471– 478.

Raven P.H. 1964. Catastrophic Selection and Edaphic Endemism. Evolution 18:336–338.

Raven P.H. 1979. A survey of reproductive biology in Onagraceae. New Zeal. J. Bot. 17:575– 593.

R Core Team. 2020. R: A language and environment for statistical computing. R Foundation for Statistical Computing, Vienna, Austria.

Roch S., Steel M. 2015. Likelihood-based tree reconstruction on a concatenation of aligned sequence data sets can be statistically inconsistent. Theor. Popul. Biol. 100:56–62.

Sayyari E., Mirarab S. 2016. Fast coalescent-based computation of local branch support from quartet frequencies. Mol. Biol. Evol. 33(7):1654–68.

Schliep K.P. 2011. phangorn: phylogenetic analysis in R. Bioinformatics. 27(4):592–593.

Schlumpberger B. O., Cocucci A. A., Moré M., Sérsic A. N., Raguso R. A. 2009. Extreme variation in floral characters and its consequences for pollinator attraction among populations of an Andean cactus. Ann. Bot. 103(9):1489–1500.

Skogen K.A., Overson R.P., Hilpman E.T., Fant J.B. 2019. Hawkmoth pollination facilitates long-distance pollen dispersal and reduces isolation across a gradient of land-use change. Ann. Missouri Bot. Gard. 104:495–511.

Smith S.A., Moore M.J., Brown J.W., Yang Y. 2015. Analysis of phylogenomic datasets reveals conflict, concordance, and gene duplications with examples from animals and plants. BMC Evol. Biol. 15:150.

Stamatakis A. 2014. Stamatakis -2014 - RAxML version 8 a tool for phylogenetic analysis and post-analysis of large phylogenies. 2010–2011.

Stanton K., Valentin C.M., Wijnen M.E., Stutstman S., Palacios J.J., Cooley A.M. 2016. Absence of postmating barriers between a selfing vs. Outcrossing chilean *Mimulus* species pair. Am. J. Bot. 103:1030–1040.

Stebbins G.L. 1970. Adaptive Radiation of Reproductive Characteristics in Angiosperms, I: Pollination Mechanisms. Annu. Rev. Ecol. Syst. 1:307–326.

Stephens J.D., Rogers W.L., Heyduk K., Cruse-Sanders J.M., Determann R.O., Glenn T.C., Malmberg R.L. 2015. Resolving phylogenetic relationships of the recently radiated carnivorous plant genus *Sarracenia* using target enrichment. Mol. Phylogenet. Evol. 85:76– 87.

Stockhouse R.E.I. 1973. Biosystematic Studies of Oenothera L. Subgenus Pachylophus. Univ. Microfilm. Ph.D. thesis. Colorado State University.

Straub S.C.K., Parks M., Weitemier K., Fishbein M., Cronn R.C., Liston A. 2012. Navigating the tip of the genomic iceberg: Next-generation sequencing for plant systematics. Am. J. Bot. 99:349–64.

Swofford, D. L. 2003. PAUP*. Phylogenetic Analysis Using Parsimony (*and Other Methods). Version 4. Sinauer Associates, Sunderland, Massachusetts.

Thomson J.D., Wilson P. 2008. Explaining Evolutionary Shifts between Bee and Hummingbird Pollination: Convergence, Divergence, and Directionality. Int. J. Plant Sci. 169:23–38.

Towner H.F., Raven P.H. 1970. A new species and some new combinations in *Calylophus* (Onagraceae). Madrono. 20: 241–245.

Towner H.F. 1977. The Biosystematics of *Calylophus* ( Onagraceae ). Ann. Missouri Bot. Gard. 64:48–120.

Tripp E.A., Manos P.S. 2008. Is floral specialization an evolutionary dead-end? Pollination system transitions in *Ruellia* (Acanthaceae). Evolution 62:1712–1737.

Turner B.L., Moore M.J. 2014. *Oenothera gayleana* (*Oenothera* sect. *Calylophus*, Onagraceae), a new gypsophile from Texas, New Mexico, and Oklahoma. Phytologia.org. 96:200–206.

Wagner W., Hoch P., Raven P. 2007. Revised Classification of the Onagraceae. Syst. Bot. Monogr. 83:1–240.

Wagner W.L. 2021. In press. In: Flora of North America Editorial Committee, eds. 1993+. Flora of North America North of Mexico, 20+ vols. New York and Oxford: Oxford University Press.

Weitemier K., Straub S.C.K., Cronn R.C., Fishbein M., Schmickl R., McDonnell A., Liston A. 2014. Hyb-Seq: Combining Target Enrichment and Genome Skimming for Plant Phylogenomics. Appl. Plant Sci. 2:1400042.

Wilson P., Wolfe A.D., Armbruster W.S., Thomson J.D. 2007. Constrained lability in floral evolution: Counting convergent origins of hummingbird pollination in Penstemon and Keckiella. New Phytol. 176:883–890.

Xi Z., Liu L., Rest J.S., Davis C.C. 2014. Coalescent versus Concatenation Methods and the Placement of *Amborella* as Sister to Water Lilies. Syst. Biol. 63:919–932.

Xu S., Schlüter P.M., Scopece G., Breitkopf H., Gross K., Cozzolino S., Schiestl F.P. 2011. Floral isolation is the main reproductive barrier among closely related sexually deceptive orchids. Evolution. 65:2606–20.

## REFERENCES

Abascal F., Zardoya R., Telford M.J. 2010. TranslatorX: Multiple alignment of nucleotide sequences guided by amino acid translations. Nucleic Acids Res. 38:7–13.

ArborBiosciences. 2016. In-Solution Sequence Capture for Targeted High-Throughput Sequencing: Version 3.02. .

Danecek P., Auton A., Abecasis G., Albers C.A., Banks E., DePristo M.A., Handsaker R.E., Lunter G., Marth G.T., Sherry S.T., McVean G., Durbin R. 2011. The variant call format and VCFtools. Bioinformatics. 27:2156–2158.

Doyle J.J. 1987. A Rapid DNA Isolation Procedure for Small Quantities of Fresh Leaf Tissue. Phytochem Bull.:11–15.

Duarte J.M., Wall P.K., Edger P.P., Landherr L.L., Ma H., Pires J.C., Leebens-Mack J., dePamphilis C.W. 2010. Identification of shared single copy nuclear genes in Arabidopsis, Populus, Vitis and Oryza and their phylogenetic utility across various taxonomic levels. BMC Evol Biol. 10:61.

Evans M.E.K., Smith S. A, Flynn R.S., Donoghue M.J. 2009. Climate, niche evolution, and diversification of the “bird-cage” evening primroses (Oenothera, sections Anogra and Kleinia). Am. Nat. 173:225–40.

G Yu, DK Smith, H Zhu, Y Guan T.L. 2017. ggtree: an R package for visualization and annotation of phylogenetic trees with their covariates and other associated data. Methods Ecol. Evol.:28–36.

Green R., Krause J., Briggs A., Rasilla Vives M., Fortea Pérez F. 2010. A draft sequence of the neandertal genome. Science (80-. ). 328:710–722.

Gutíerrez S.C., Martínez J.M.S., Gabaldón T. 2009. TrimAl : a tool for automatic alignment trimming. Bioinformatics.:1–2.

Illumina. 2016. BaseSpace Sequence Hub. Available from https://basespace.illumina.com/.

Johnson M.G., Gardner E.M., Liu Y., Medina R., Goffinet B., Shaw A.J., Zerega N.J.C., Wickett N.J. 2016. HybPiper: Extracting Coding Sequence and Introns for Phylogenetics from High-Throughput Sequencing Reads Using Target Enrichment. Appl. Plant Sci. 4:1600016.

Katoh K., Misawa K., Kuma K., Miyata T. 2002. MAFFT: a novel method for rapid multiple sequence alignment based on fast Fourier transform. Nucleic Acids Res. 30:3059–3066.

Li H., Durbin R. 2009. Fast and accurate short read alignment with Burrows-Wheeler transform. Bioinformatics. 25:1754–1760.

Miller M.A., Pfeiffer W., Schwartz T. 2010. Creating the CIPRES Science Gateway for inference of large phylogenetic trees. 2010 Gatew. Comput. Environ. Work. GCE 2010.

Minh B.Q., Hahn M.W., Lanfear R. 2018. New methods to calculate concordance factors for phylogenomic datasets. bioRxiv.:487801.

Paradis E., Claude J., Strimmer K. 2004. APE: Analyses of phylogenetics and evolution in R language. Bioinformatics. 20:289–290.

Poplin R., Ruano-Rubio V., DePristo M.A., Fennell T.J., Carneiro M.O., Auwera G.A. Van der, Kling D.E., Gauthier L.D., Levy-Moonshine A., Roazen D., Shakir K., Thibault J., Chandran S., Whelan C., Lek M., Gabriel S., Daly M.J., Neale B., MacArthur D.G., Banks E. 2017. Scaling accurate genetic variant discovery to tens of thousands of samples. bioRxiv.:201178.

Purcell S., Neale B., Todd-Brown K., Thomas L., Ferreira M.A.R., Bender D., Maller J., Sklar P., De Bakker P.I.W., Daly M.J., Sham P.C. 2007. PLINK: A tool set for whole-genome association and population-based linkage analyses. Am. J. Hum. Genet. 81:559–575.

R Core Team. 2018. R: A language and environment for statistical computing.

Sayyari E., Mirarab S. 2016. Fast coalescent-based computation of local branch support from quartet frequencies. Mol. Biol. Evol.:msw079.

Schliep K.P. 2011. phangorn: Phylogenetic analysis in R. Bioinformatics. 27:592–593.

Sharma P., Purohit S.D. 2012. An improved method of DNA isolation from polysaccharide rich leaves of Boswellia serrata Roxb. Indian J. Biotechnol. 11:67–71.

Stamatakis A. 2014. Stamatakis - 2014 - RAxML version 8 a tool for phylogenetic analysis and post-analysis of large phylogenies. :2010–2011.

Sukumaran J. and M.T.H. 2010. DendroPy: A Python library for phylogenetic computing. Bioinformatics.:1569–1571.

Suyama M., Torrents D., Bork P. 2006. PAL2NAL: Robust conversion of protein sequence alignments into the corresponding codon alignments. Nucleic Acids Res. 34:609–612.

Towner H.F.. 1977. The Biosystematics of Calylophus ( Onagraceae ). Ann. Missouri Bot. Gard. 64:48–120.

Wickham H. 2016. ggplot2: Elegant Graphics for Data Analysis. Springer-Verlag.

Zhang C., Rabiee M., Sayyari E., Mirarab S. 2018. ASTRAL-III: polynomial time species tree reconstruction from partially resolved gene trees. BMC Bioinformatics. 19:153.

